# Evolution at experimental epidemic fronts speeds up parasite spread

**DOI:** 10.64898/2026.01.06.697932

**Authors:** Manasven Raina, Margherita Allotta, Julien Lombard, Jhelam N. Deshpande, Charlène Sanchez, Claire Gougat-Barbera, Leandro Gammuto, Giulio Petroni, Oliver Kaltz, Giacomo Zilio

## Abstract

Spreading epidemics can foster the rapid emergence of novel parasite variants. Similar to range expansions, parasites at the front of an epidemic wave may evolve traits facilitating their spatial spread. Typically, higher transmissibility and virulence are expected to evolve at the epidemic front, but predictions change if virulence reduces dispersal of infected hosts. We investigated the feedbacks between epidemiology and evolution in interconnected microcosms of the ciliate *Paramecium caudatum* and its bacterial parasite *Holospora undulata*. First, two long-term treatments mimicked the moving front of epidemic waves, with and without the natural dispersal of infected hosts. Phenotypic trait assays revealed an eco-to-evo feedback: front parasites (*i.e.*, travelling with infected hosts) were less virulent and interfered less with host dispersal, but also showed higher infectivity, compared to parasites from the control treatment. Whole-genome resequencing and analyses of functional genomic diversity corroborated such phenotypic divergence. Second, measurement of the spread of evolved parasites in linear landscapes demonstrated an evo-to-eco feedback: front parasites produced stronger infection outbreaks and faster spreading epidemic waves than did control parasites. A simulation model assessed the relative importance of the different parasite traits in determining infectious spread over a broad parameter range. It suggests that, if dispersal is generally low, higher infectivity alone can produce the observed differences in epidemic spread speed. Our study illustrates how the need to travel with the host shapes eco-evolutionary feedbacks at parasite invasion fronts, and it highlights the importance of considering concomitant evolutionary change when predicting epidemic speed.

**Significance Statement:** When predicting disease spread across spatial scales it is fundamental to consider host dispersal, but also the concomitant evolution of parasite traits such as infectivity and virulence. We tested how dispersal constraints affect parasite evolution and epidemic spread in an aquatic model system, by combining experimental evolution, genetic analysis and mathematical modelling. Our results suggest that epidemics can be accelerated by the evolution of highly infectious, but not necessarily more virulent parasites, in contrast to conventional views. This illustrates the important role of eco-evolutionary feedback loops can play in shaping the spread of emerging or geographically expanding parasites.

## Introduction

Infectious diseases represent a growing threat with climate change, habitat disruption or increased host mobility allowing their spread over large geographic scales (1–6). In this context, the evolutionary potential of many parasites is of particular concern. Rapid evolution during epidemics may not only add uncertainty regarding the harmfulness of disease (virulence), but also impact predictions of the speed of epidemic spread. Yet, to date, it is not evident whether parasite evolution follows general rules (7). When disease is spreading, it is conceivable that selection rapidly picks up any kind of adaptation enhancing local host-to-host transmission or facilitating dispersal to new infection sites, in particular when populations are spatially structured (8–10). Understanding the eco-evolutionary feedbacks produced by such adaptations is essential to fundamental research (10, 11), but might also improve epidemiological forecast models or the evaluation of emerging diseases (12–14).

Empirical studies combining epidemiological and genomic data show that the epidemic spread of parasites across populations and continents often coincides with the divergence into distinct genetic lineages (15–17). This can involve the evolution of fitness-relevant traits. For example, the global succession of viral lineages of SARS-CoV-2 was associated with increased transmissibility (18), similar to reports for other human viruses (19). Along the same lines, natural systems also display spatial variation in life-history, sensitivity to host defence or virulence, emerging in the context of biological invasions or for migratory host species (20–24).

Theoretical models have investigated evolutionary processes in spatially expanding epidemics (11, 13, 25). They show that the transient nature of epidemics yields outcomes distinct from classic equilibrium models, due to eco-evolutionary feedbacks (26, 27). For instance, Griette *et al.*, (30) predict the evolution of increased parasite investment in transmission and thus higher virulence at the front of epidemic waves, where susceptible hosts are abundant. Behind the front, lower virulence is favoured due to host depletion and kin competition (11, 27–30). However, the opposite is predicted if virulence trades off with host dispersal, with wave fronts harbouring fast-advancing, but less virulent variants, escaping higher-fitness variants from behind the front via dispersal (31). These different models highlight two important aspects. First, the front of an epidemic wave represents specific ecological conditions, to which parasites adapt as they move along, with constant spatial sorting favouring variants that are best at getting to and exploiting new patches of hosts (eco-to-evo feedback). Second, parasite adaptation at the epidemic front produces an evo-to-eco feedback, with evolution accelerating the rate of spatial epidemic spread. This is similar to the auto-fuelling dynamics of range expansions, where the evolution of dispersal syndromes the expansion front speeds up spatial progression (32, 33).

Verifying these predictions from empirical data remains a challenge because it is difficult to assess whether these parasite changes and emerge of variants are driven by spatial selection at the epidemic front, as envisaged by the theory, or by other factors, such as environmental variation, differences in host responses, or simply genetic drift. Identifying evo-to-eco feedbacks is even more difficult, because adaptive evolutionary change needs to be linked with spatio-temporal data of epidemic spread. This is complicated for data from historical records (34), but also for contemporary phylodynamic approaches, estimating epidemic spread speed from (near-)neutral genetic variation (6, 15, 35).

Alternatively, theory can be tested in experimental host-parasite systems, under controlled conditions in replicated micro- or mesocosms (36). Spatial aspects have been investigated in several studies (reviewed in (10)). For example, viruses of bacteria and insects evolved higher virulence under experimentally enhanced spatial mixing (37–39) or at the border of phage plaques on bacterial lawns (40), consistent with the prediction that access to susceptible hosts is a main evolutionary driver. In contrast, Nørgaard *et al.* (43) showed that continued spatial spread of a bacterial parasite (*Holospora undulata*) with its dispersing ciliate host (*Paramecium caudatum*) favoured less virulent variants, revealing an evolutionary virulence-dispersal trade-off (31). Together with other dispersal-related experiments (10), these examples demonstrate that conditions of spatial spread can indeed produce eco-to-evo feedbacks. However, still few studies have employed explicit meta-population settings with natural dispersal, and practically none have quantified the resulting evo-to-eco feedback, that is, by how much parasite evolution accelerates spreading epidemics (40).

We studied both parts of the eco-evo feedback loop, combining experimental evolution, whole-genome sequencing and epidemiological modelling. First, we assessed the eco-to-evo feedback, by letting the above-mentioned bacterial parasite *H. undulata* evolve under conditions that mimicked spatially spreading epidemics, accomplished via the dispersal of infected hosts (*P. caudatum).* Evolved parasites and those from a corresponding control treatment were compared for phenotypic differentiation in key fitness traits (virulence, dispersal, and transmission), and their genomes sequenced. Second, to assess the evo-to-eco feedback, we let the two types of parasites produce epidemic waves in microcosm landscapes, and compared spread speed and outbreak intensities. We also recreated epidemic waves with numerical simulations, assessing the impact of parasite trait variation across wider parameter space.

## Results

### Eco-to-evo feedback: Experimental parasite evolution at an epidemic front

In a long-term experiment, we let infected populations regularly colonise a new “front patch”, thereby simulating spatial progression. In the wave-front treatment (front from now onwards), colonisation occurred through the active dispersal of infected hosts between interconnected tubes (**Fig. S1**). By making dispersal with infected hosts obligate (the parasite itself is immobile), this treatment imposed host-mediated spatial selection on the parasite (see also (41)). In the control treatment (control from now onwards), we relaxed dispersal-related constraints by manually transferring arbitrarily picked samples of hosts and parasites to the new patch. After 9 months (39 dispersal events), we extracted parasites from each of 6 front and control lines and assayed them for several traits on naïve hosts.

#### Parasite transmission capacity

In a natural transmission test, 5 infected hosts were introduced into groups of 40 uninfected individuals. Over 5 days, parasites from the front treatment produced more secondary infections (mean = 22, 95% CI = [15.2; 30.88]) than control parasites (14.69, CI [9.89; 20.99]), indicating a higher transmission capacity of front parasites (χ^2^_1_ = 3.94, p = 0.04; **Fig. 1A**, **Table S1**).

**Figure 1.**
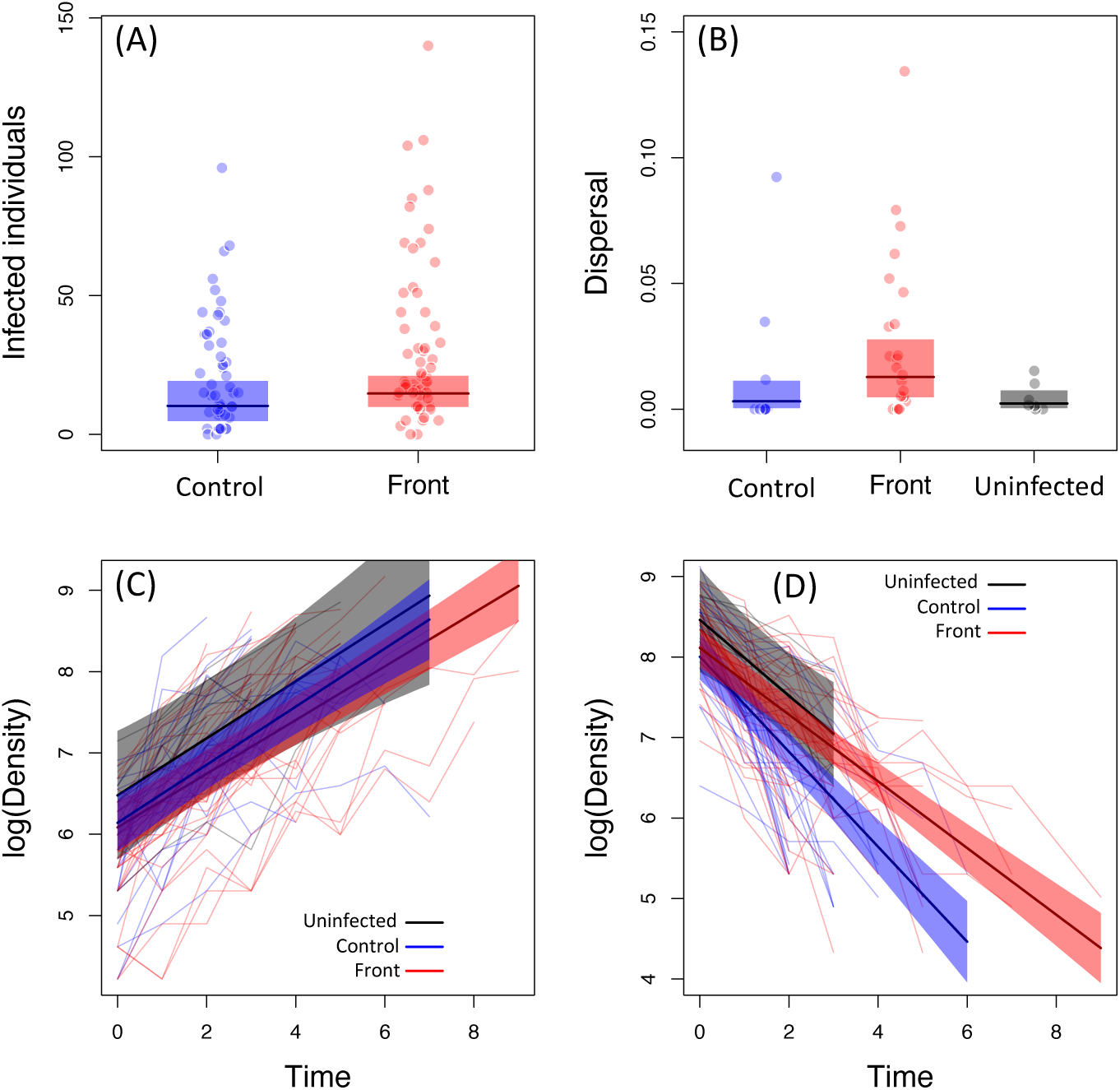
Transmission, dispersal and virulence of parasite evolved in the control (blue) or at the front (red) of epidemic waves measured on naïve host; naïve uninfected control replicates are represented in black. Each point and line represent and individual replicate, panels the 95% confidence intervals of the model predictions, and thick lines the mean model prediction. (A) Transmission. Number of infected individuals (secondary infections) produced over 5 days by placing 5 infected with 40 uninfected individuals. (B) Dispersal. The proportion of dispersing infected hosts during 3h in 2-patch dispersal system. In black the dispersal of uninfected individuals. (C, D) Virulence. Exponential growth (C) and decline phase (D) obtained from growth assay as logarithm of population density over time. Densities were sampled at least once per day (see main text). Time was translated to the same starting point.

#### Virulence

In a population growth assay, infection had no significant affect during the exponential growth (evolutionary treatment*time: (χ^2^_2_ = 0.33, p > 0.8, **Fig 1C**, **Table S2**), but differences appeared during the decline phase (χ^2^_2_ = 8.62, p = 0.01; **Fig. 1D**, **Table S3**). Infection with control parasites accelerated population decline (slope = -0.59, 95% CI [-0.69, -0.48]), whereas front parasites was less virulent causing rates comparable (-0.41, CI [-0.47, -0.35]) to those of uninfected cultures (-0.47, CI [-0.72, -0.21]).

#### Dispersal

We assayed the dispersal of infected hosts in two-patch arenas (**Fig. S1**). A significant treatment effect was detected (χ^2^_2_ = 6.54, p = 0.03; **Fig. 1B**, **Table S4**). Front parasites tended to stimulate host dispersal, with over 80% of replicate cultures infected with front parasites had non-zero dispersal. Mean dispersal rates were 4-5 times higher (0.012; CI [0.004; 0.027]) than control parasites (0.003; CI [0.0004; 0.011]) or uninfected hosts (0.002; CI [0.0003; 0.007]).

### Eco-to-evo feedback: Genomic analysis

Whole-genome sequencing of evolved parasites sampled after 6 months of treatment revealed 104–486 single-nucleotide-polymorphism (SNP) variants per evolved line (**Table S5**). In a phylogenetic analysis, front and control lines formed distinct clusters, with the former being closer to the ancestral genome (**Fig. 2A**).

**Figure 2.**
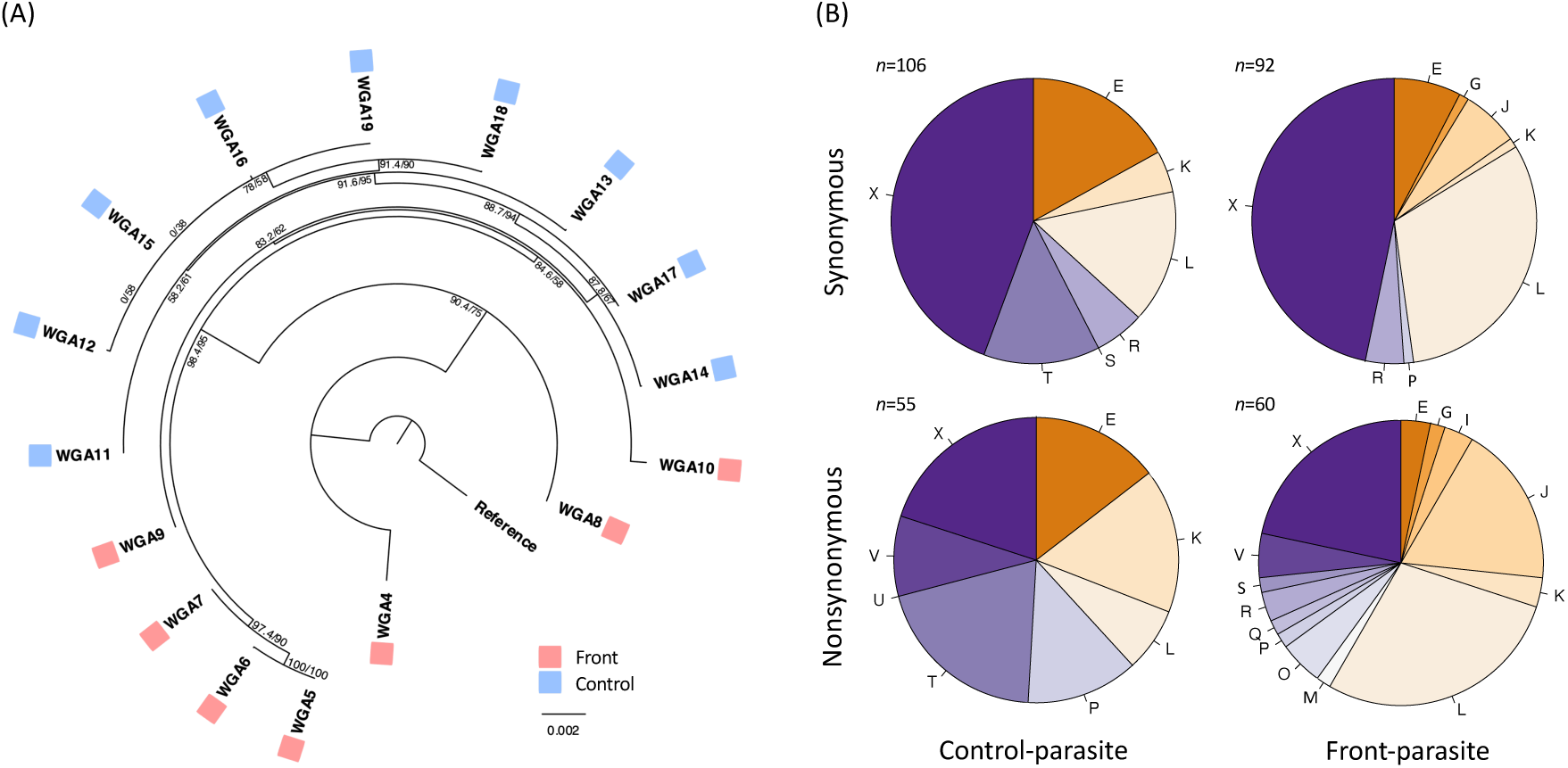
(A) Phylogenetic divergence of front parasites (7 evolved lines) and control parasites (9 lines) estimated from whole-genome resequencing data using a maximum-likelihood technique. Numbers on nodes indicate SH-aLRT index and Ultra-Fast Bootstrap. (B) Distribution of synonymous SNP variants (top) and non-synonymous variants (bottom) in genes from 21 classes of Clusters of Orthologous Groups (COGs), for front and control parasites.

To investigate signatures of selection, we analysed the frequency of synonymous (S) and non-synonymous (N) variants of genes with generally known bacterial functions, defined by 21 Clusters of Orthologous Groups (COGs, **Table S6**). Overall, there was no evidence for a genome-wide difference in dN/dS ratios between front and control parasites (χ^2^_1_ = 2.69, p = 0.10; **Table S7**). However, we did find differences when taking into account COG diversity. For synonymous variants, front and control showed similar frequency distributions over a total of 9 COGs (COG*treatment: χ^2^_1_ = 19.64, p = 0.48; **Fig. 2B** top row; **Fig. S2A**; **Table S8**). In contrast, front parasites had a greater non-synonymous variant diversity, with 60 variants (from 7 parasite lines) distributed across 14 COGs, whereas the 55 control variants (from 9 lines) were confined to only 7 COGs (COG*treatment: χ^2^_1_ = 44.67, p = 0.001; **Fig. 2B** bottom row; **Fig. S2B**; **Table S9**). Certain COGs harboured exclusively non-synonymous variants (**Fig. S2B**), all of which occurred in front parasites, suggesting possible treatment-specific responses to selection. These COGs are associated with lipid metabolism (I), cell wall biogenesis (M), protein turnover (O), and secondary metabolite biosynthesis (Q) in bacteria, although their functions in *Holospora* have not been described yet.

### Evo-to-eco feedback: evolutionary divergence and epidemic spread in landscapes

To assess the impact of evolutionary differentiation on epidemiology, we allowed evolved parasites to spread in linear landscapes of interconnected tubes (**Fig. 3**). On one edge (patch 1, **Fig. 3**), a population of naïve *Paramecium* received a small number of hosts infected with a given parasite (initial prevalence 10%). Regular opening of connections then enabled dispersing *Paramecium* to colonise downstream tubes over 4 weeks (16 openings).

**Figure 3.**
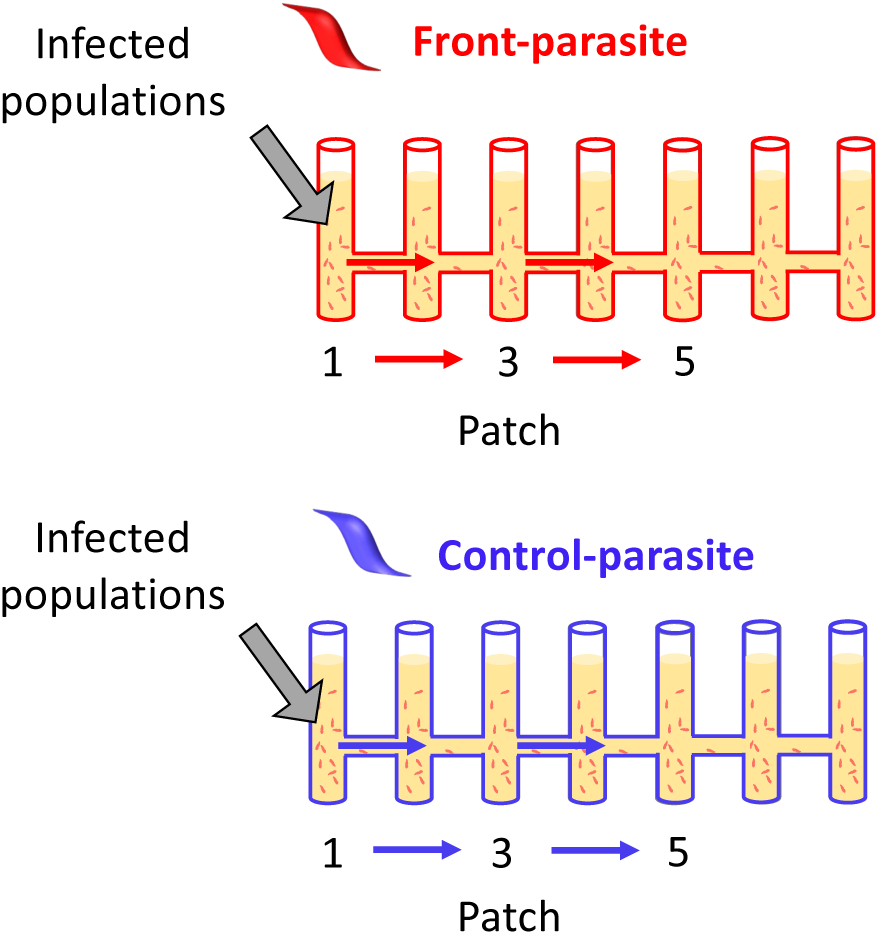
Schematic representation of the linear 7-patch landscapes used for the epidemiological assays. Naïve hosts infected with parasites from the front (red) or control (blue) treatments were introduced in patch 1, which was already filled with naïve uninfected hosts. All other patches were empty at the start of the experiment. Connections to downstream patches were opened 3 times per week for 3h, allowing the free dispersal of infected and uninfected hosts and colonisation of new patches (red and blue arrows).

Typically, uninfected hosts arrived one to several time points before the parasite (**Table S10**; **Fig. 4A, B**). Hence, infections spread into already populated patches, producing wave-like epidemics, with outbreaks shifted in space and time (**Fig. 5**). Because in many landscapes (13/23) infection had not reached the outermost patch 7 by the end of the experiment, we only report on the dynamics in focal patches 1, 3 and 5.

**Figure 4.**
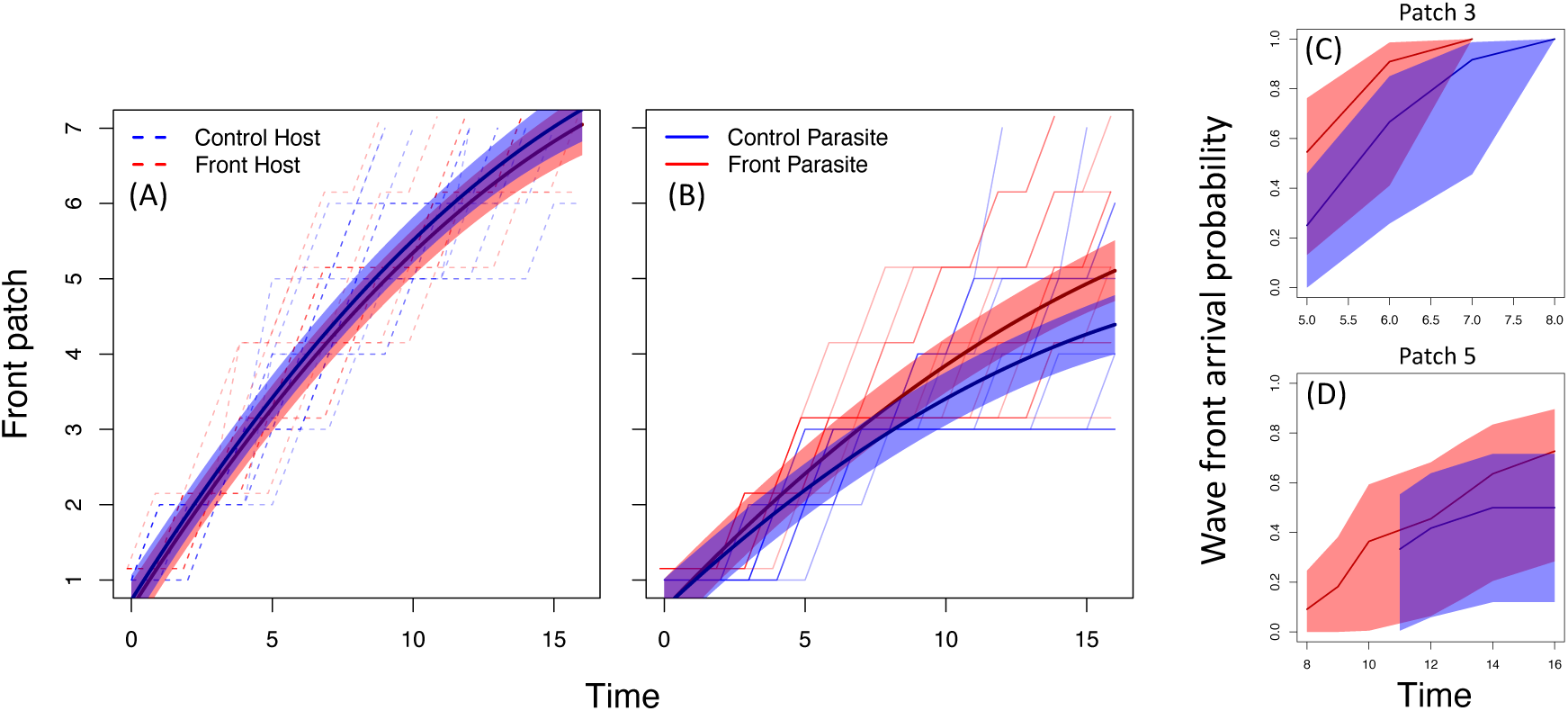
Advancing front position and speed of the epidemic wave during the time of the experiment (opening events). (A) Front patch position of uninfected hosts within infected populations with control (blue) or front (red) evolved parasite, and (B) front patch position of infected hosts with control (blue) or front (red) evolved parasites. Each line is an individual replicate, panels show the 95 % confidence intervals of the model predictions. Thick lines are mean model prediction. (C) Epidemic wave front as parasite time of arrival from patch 1 to patch 3 over opening events, and (D) from patch 3 to patch 5 over opening events. Thick lines and panels are the proportion of replicates arriving at certain time points with 95% confidence intervals.

**Figure 5.**
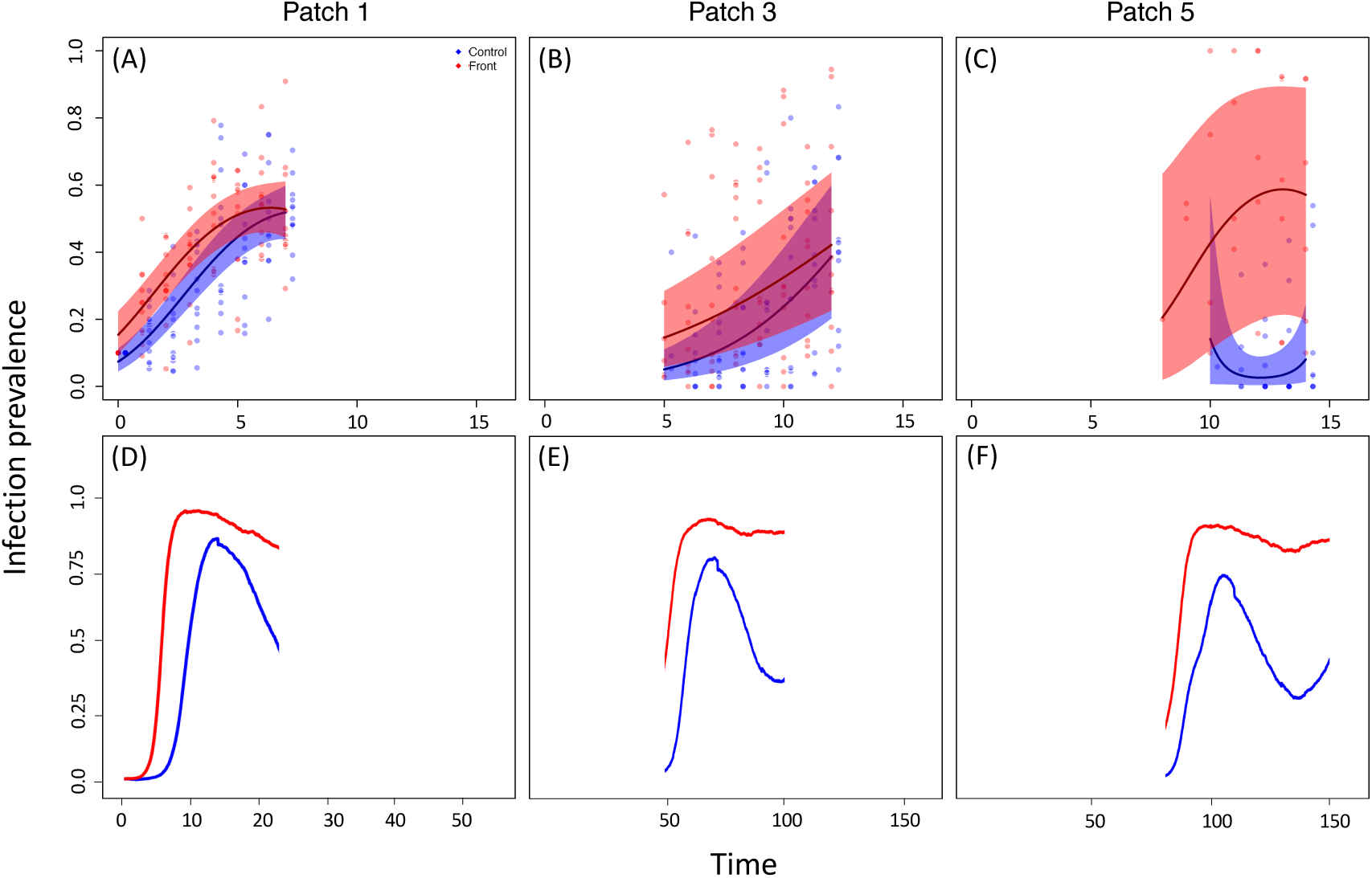
Infection prevalence and number of infected individuals over time (opening events) in patch 1, 3 and 5 of the epidemiological assays and model simulations. (A-C) Epidemiological assays. Control and front parasites are in blue and red respectively. Each point represents an individual replicate measurement, panels the 95 % confidence intervals of the model predictions, and thick lines the mean model prediction. (D-F) Model simulations. Infection prevalence dynamics obtain from single-run simulations. Curves are obtained by averaging replicates of a given parameter combination. The blue curve (control) corresponds to a parasite with λ = 0.25, d = 0.4, α = 0.64, β = 7.9.10−5, while the red (front) corresponds to a parasite with λ = 0.94, d = 0.4, α = 0.12, β = 1.5.10−4. Others parameters values were fixed and set to values in Table S13.1. Infection prevalences for patch 7 are not shown due to few replicates establishing (see Material and Methods and Fig. 4B).

We found that, first, wave-front parasites produced stronger local outbreaks than did control parasites, with prevalences reaching 60–80%. This pattern established over the first four time points in patch 1 (evolution treatment*time: χ^2^_1_ = 6.84, p = 0.008; **Fig. 5A**; **Table S11**), and was repeated in downstream focal patches 3 (χ^2^_1_ = 5.94, p = 0.01; **Fig. 5B**; **Table S12**) and 5 (χ^2^_1_ = 4.97, p = 0.02; **Fig. 5C**; **Table S13**).

Second, wave-front parasites showed faster spatial spread. In patch 3, infection appeared in >50% (6/11) of wave-front replicates by time point 5, compared to only 25% (3/12) for control replicates. Similarly, in patch 5, the first wave-front parasites arrived two time points earlier than control parasites (**Fig. 5C**). Accordingly, analysis of parasite position as a function of time found steeper slopes for wave-front parasites (evolution treatment*time: χ^2^_1_ = 12.72, p = <0.001; **Fig. 4B**; **Table S14**), indicating higher spread speed. The same trend was found in a complementary analysis, showing significantly shorter time to arrival of wave-front parasite in downstream patches (treatment: χ^2^_1_ = 3.98, p =0.04; **Fig. 4C-D**; **Table S15**). By the end of the experiment, prevalences had generally settled to lower levels, but wave-front parasites still showed higher landscape-wide infection prevalence, namely in the more recently colonised downstream patches 4 and 5 evolutionary (treatment * patch: χ^2^_1_ = 19.91, p = <0.001; **Fig. S2**; **Table S16**).

### Epidemiological model: parasite trait variation and epidemic waves

We created a simulation model, integrating key features of our experimental setup (**Table S17**) and producing similar epidemic waves (Fig. 5D-F). We varied four parameters (host dispersal, virulence, horizontal transmission, and parasite-mediated dispersal modification; **Table S17**) and inspected their impact on epidemiological dynamics, using sensitivity analysis (**Table S18**).

First, local disease spread (= epidemic peak size) was mainly driven by variation in virulence and transmission (13% and 11.7% of variation explained, respectively; **Table S18**). Because the simulation model assumes no a priori relationship between the two parameters, the strongest outbreaks arose for very high transmission and very low virulence (**Fig. 6**). However, virulence caused negative demographic feedbacks, resulting in non-linear trends. For high levels of transmission (β, **Fig. 6A**), even moderate increases in virulence caused a decline in epidemic peak size, whereas for low transmission, the decline in peak size was stretched out over a larger virulence range (**Fig. 6A**).

**Figure 6.**
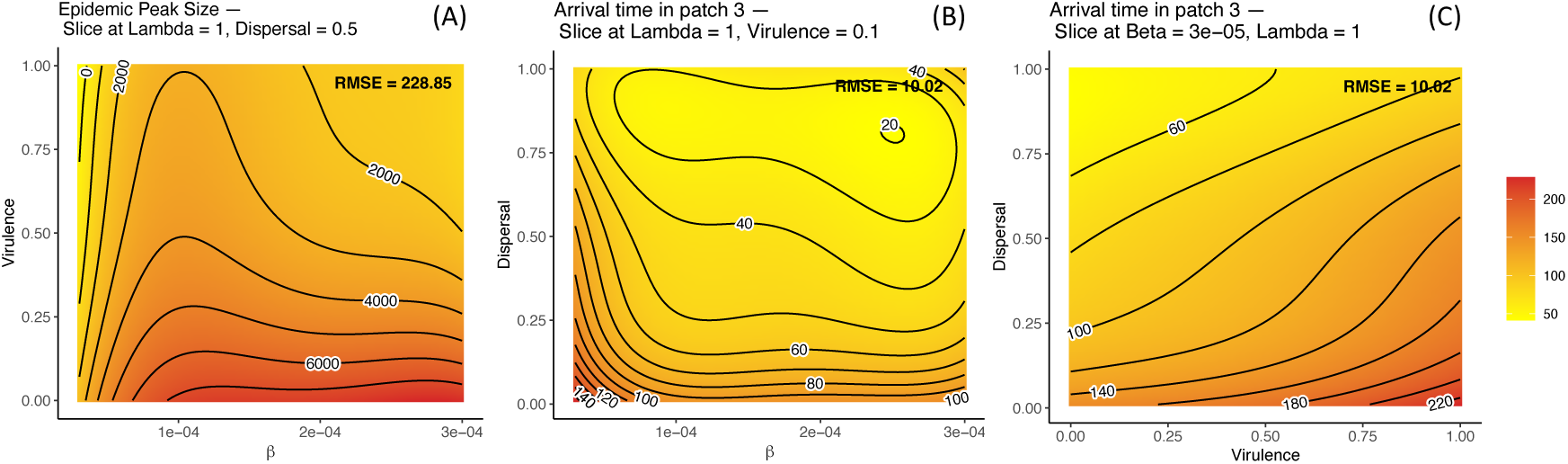
Heatmaps obtained from the response-surface on our set of simulated data. (A) Average local epidemic peak size as a function of the virulence and horizontal transmission rate. (B) Arrival times in patch 3 with respect to the horizontal transmission and dispersal rates. (C) Arrival times in patch 3 with respect to the dispersal rate and virulence. The colour gradient indicates in yellow lower epidemic peak size (A) or earlier arrival times (B, C), and in red higher epidemic peak size (A) or later arrival times (B, C). Constant parameters were set according to values in Table S13.1.

Second, the speed of spatial parasite spread (= time to arrival in patch 3) was most dependent on host dispersal, due to the fact that parasites travel with infected hosts (15.6% variation explained, Table S18). Transmission and virulence also affected spatial spread (4% and 7.2%, respectively), again indirectly, through a demographic feedback. Both high transmission and low virulence increase infected host density, thereby increasing the probability of infected individuals moving to a new patch (**Fig. 6B, C**; see also equation 3 in Material & Methods). In contrast, direct parasite impact on host dispersal (dispersal modulator, λ) had little effect (**Table S18**), meaning that status-dependent host dispersal contributes little to epidemic spread in our simulations.

## Discussion

Spatially spreading epidemics resemble biological invasions or range expansions. These can create characteristic eco-evolutionary feedbacks where selection for high dispersal and fast growth at the front accelerates expansion (32, 33). Our study of experimentally spreading epidemic waves revealed similar feedbacks. Analogous to dispersal syndromes in range expansions (42), multi-trait associations emerged in parasites from the front treatment, enhancing transmission and facilitating dispersal. These adaptations likely explain the faster epidemic waves in the microcosm landscapes.

### Eco-to-evo feedback: the evolution of parasite traits at epidemic fronts

In our front treatment, spatial spread of the parasite depended on dispersal of infected hosts. Osnas *et al.* (33) showed that this constraint favours lower virulence at epidemic fronts. Consistent with this, front parasites were less virulent and even tended to increase host dispersal compared to parasites from the control (**Fig. 1B**). Our type of experimental design is known to impose strong selection for increased dispersal in the host (43–45), and clearly the parasite, obligately travelling with host, is exposed to the same selection pressure. As host exploitation often reduces mobility (46, 47), lower virulence is therefore a plausible route to enhance dispersal, as observed here and in a previous study (41). Virulence–dispersal trade-offs have also been reported for a bacterial parasite of water flees (48) and for a protozoan parasite of natural migratory butterfly populations (49), suggesting that host dispersal manipulation could be an evolving trait (50, 51).

Front parasites were also more infectious than control parasites (**Fig. 1A**). Increased transmissibility often evolves in spreading diseases (18, 19, 52), presumably because parasites at wave fronts encounter abundant susceptible hosts (28). Our treatments did not explicitly manipulate host availability. However, because infection reduces host dispersal (46, 47), relatively more uninfected hosts may arrive in new front patches in the front than in the control treatment bypassing natural dispersal. This is consistent with the lower prevalences observed early the experiment in the front treatment (**Fig. S4**), which may have favoured greater investment in horizontal transmission.

Theory often links higher transmissibility to greater production of transmission stages and thus higher virulence (27, 28). However, evolved front parasites were less virulent than control parasites, and showed no significant difference in spore production (**Fig. S5**; **Table S19**), suggesting transmission investment is not strongly resource-limited. Increased infectivity has also been reported for a parasitic lungworm from expanding populations of the cane toad (*Rhinealla marina*) in Australia (21, 24), where, as in our study, greater infectivity is not associated with higher virulence (53). We therefore hypothesise that selection can act on the quality, rather than quantity, of transmission stages. Studies on microparasites support this idea and highlight the need to investigate this evolutionary pathway more closely (54).

### Molecular analysis corroborates phenotypic divergence, but targets of selection remain unclear

Genomes of front and control parasites formed distinct clusters after 30 dispersal events, indicating rapid evolutionary divergence. Front parasites appeared genetically closer to the ancestor than control parasites. This is unexpected, as spatial selection was predicted to operate in the front treatment, thereby drive divergence. As discussed above, unintended additional selective forces may have acted in our treatments. However, we found front genomes more diverged from the ancestor when examining non-neutral variants in genes with annotated bacterial functions (**Fig. 2B**). This may reflect treatment-specific evolution in genes associated with metabolic function, within-cell trafficking or signalling (**Table S6**), though their precise role in *Holospora* is unknown. We also note that this putative signal of selection is limited, involving only a relatively small number of variants from certain front lineages. Future sequencing efforts, with improved coverage and less fragmented genome assembly, will allow more comprehensive annotation of the *Holospora* genome and help identify targets of selection.

### Evo-to-eco feedback: faster epidemic spread due to parasite evolution

The first part of our work joins a small body of “eco-to-evo” studies showing how spatial structure shapes parasite evolution (8, 10). The second part demonstrates the reverse “evo-to-eco” feedback on epidemic spread: Parasites evolved in the front treatment produced substantially stronger and faster spreading outbreaks than control parasites.

Stronger local outbreaks are readily explained by the higher infectivity of front parasites. The time scale of outbreaks (5-10 days) matches a scenario where incoming infected hosts produce many secondary infections, as seen in the infectivity assay (**Fig. 1A**). Identifying the traits driving landscape-level spread is more difficult. Following Osnas *et al.* (33) and extrapolating from our trait assays (see also (41)), both lower virulence and higher infected-host dispersal of ront parasites could explain their accelerated spread. Yet, there were no marked differences in their effects on host density in the landscapes (**Fig. S6**, **Table S20**), and dispersal was not formally quantified, offering limited direct support for this explanation. A simple (non-exclusive) alternative is the higher transmission of front parasites: by infecting more hosts, these parasites would increase their probability of being dispersed to a new patch, all else being equal. This is pertinent, given the small infected population sizes and low colonization rates (<1 patch per dispersal event; **Fig. 4B**).

The simulation model also highlights the role of demographic stochasticity. It finds that both highly transmissible and avirulent variants are the most effective epidemic spreaders (**Fig. 6**; **S18**). At the local scale this expected, as this trait combination generates large numbers of infected individuals during outbreaks. As argued above, this can also explain higher spatial spread rate (**Fig. 6**), by increasing the chance that infected hosts disperse to new patches. Consistent with this, when dispersal is low, local infected density predicts parasite arrival time in the next downstream patch in the simulations (**Fig. S7**).

The combination of low virulence and high transmissibility emerged naturally in our experiment and was also most successful in the simulations, yet it would be prohibited in traditional models assuming a fixed virulence-transmission trade-off (Anderson & May 1982). Our model does not impose such a constraint, but still shows that virulence generates negative demographic feedbacks. Already at intermediate transmission levels, virulence reduces epidemic peak size (**Fig. 6**). This local effect propagates to the spatial scale, producing a demographic virulence-dispersal trade-off. Thus, low-virulence variants may be favoured at the front of spreading epidemics, even without an explicit virulence-movement trade-off (31), provided population sizes are small enough for stochastic effects to matter.

Direct parasite modification of host dispersal (*λ*) had only minor influence in the simulations. Because spatial parasite spread depends on the product of *λ* and host dispersal, effects of the state-dependent component may have been swamped by that of the dispersal component. Indeed, host dispersal rate was the main determinant of spread speed, explaining most variation in spatial parasite spread speed (**Table S18**). This is a reminder that knowledge of basic host biology (here, dispersal) can matter as much as detailed insight in parasite trait variation.

### Concluding remarks

Host mobility and dispersal determine epidemic spread across larger spatial scales (55, 56). This study illustrates how dependence on host dispersal shapes eco-evolutionary feedbacks at parasite invasion fronts. It suggests that epidemics can be accelerated by highly infectious but not necessarily more virulent variants, challenging conventional views of parasite evolution. Such deviations from expected trajectories may be characteristic of emerging or expanding parasites (7), which calls for eco-evolutionary models that capture the relevant stages of the infection life cycle, and include realistic representations of how parasites (and hosts) disperse (57, 58). To infer feedback loops in naturally progressing epidemics will require data linking epidemic spread speed with changes in parasite fitness.

## Materials and Methods

### Study system

We maintain *Paramecium caudatum* cultures at 23 °C in a lettuce medium, supplemented with the bacterium *Serratia marcescens* (59). Infection with the gram-negative bacterial parasite *Holospora undulata* (60) occurs via ingestion of immobile infectious forms. The parasite multiplies in the micronucleus and is vertically transmitted in mitotically dividing hosts, with infectious forms released after host division or death. Infection reduces *Paramecium* survival and division (61). From small samples of culture, we measure population density by counting individuals under a dissecting microscope and infection prevalence by determining infection status with staining and phase-contrast microscopy (62).

### Long-term experiment

We infected a culture of *P. caudatum* (strain KS3, (63)) with mixed inocula from *H. undulata* stock cultures (strain 255 (64)). From 50 arbitrarily picked cells, we then established a new infected culture for the long-term experiment. The front treatment mimicked epidemic expansion in two-patch systems (30 mL tubes connected by 5 cm silicon tubes (41)). We placed 20 ml of culture (∼2000 individuals) in one tube and allowed dispersal into the other tube for 3h (**Fig. S1**). Dispersers were then transferred to 100 ml of lettuce medium in 250 ml bottles. During the following week, host population growth and infection dynamics unfolded naturally, until the next dispersal episode. The control treatment followed the same protocol, but bypassed the natural dispersal step, with arbitrarily picked samples being manually pipetted to new bottles, at densities matching the front treatment. We initiated nine replicate selection lines per treatment, though two front lines were lost early in the experiment.

### Parasite trait assays

After 39 dispersal–growth cycles, we extracted infectious forms from each of 6 arbitrarily picked infected selection lines per treatment and used these inocula to infect naïve KS3 host cultures from background stocks. For each parasite selection line, we then established multiple, independent infected assay cultures, each derived from a single infected individual. These assay cultures were used in the tests below, employing up to 9 infected assay cultures per parasite line and 4–5 naïve uninfected KS3 assay cultures.

### Transmissibility

In 2-ml tubes, we mixed 40 uninfected individuals with 5 hosts infected with a given parasite and carrying many horizontal stages (established infections). Epidemiological dynamics then unfolded naturally. After 5 days, we quantified the number of newly infected hosts (secondary infections). We ran 6–10 tests per parasite line (104 replicates).

### Virulence

In population growth tests, we placed 100 hosts infected with a given parasite in 20 ml of medium (in 50-ml tubes). Densities were tracked for 3 weeks (28 time points), covering population growth, equilibrium, and decline. We set up 69 infected replicates (≥ 4 replicates per parasite line) and 6 uninfected replicates. Six infected replicates failed to develop and were discarded.

### Dispersal

In two-patch systems (interconnected 15 ml tubes; **Fig. S1**) we introduced 8 ml of infected culture (∼1000 hosts) into one tube and allowed dispersal to the other tube for 3 h. Density measurements in both tubes gave the proportion of dispersers. We tested 5 front and 5 control parasite lines (≤5 replicates each), plus 7 uninfected replicates (38 tests total).

### Genome sequencing

Parasite DNA was extracted from samples of the ancestral population and from 16 long-term lines after 6 months of evolution (NucleoSpin Plant II kit; Macherey-Nagel, Germany). After Whole Genome Amplification (REPLI-g Single Cell Kit, QIAGEN, Germany), Illumina sequencing generated 150 bp paired-end reads (Genewiz/Azenta Life Sciences, Leipzig, Germany). We checked read quality with FastQC, and trimmed low-quality bases and Illumina adapters with Trimmomatic (65). Ancestral reads were mapped to the *H. undulata* reference genome (RefSeq: GCF_000388175.3), assembled with SPAdes v3.36 (66), and annotated using PROKKA (67). Variant calling was performed with SNIPPY (68), comparing evolved and ancestral genomes. We used IQ-TREE 2 (69) for phylogenetic tree construction, aligning evolved and ancestral sequences for 29717 genomic sites, of which 266 were informative. The K2P+I substitution model was used, and branch support assessed from 1000 ultrafast bootstrap replicates and 1000 SH-aLRT replicates (83) We further analysed the predicted proteins, assigning them to different classes of Clusters of Orthologous Groups (COGs) using the NCBI COG pipeline and the COG database version 2014 (70). We also quantified the number of non-synonymous variants (missense and frameshift mutations) and synonymous variants for these genes.

### Landscape assay

Epidemic waves were studied in linear landscapes of 7 interconnected 30-ml tubes (Fig. 3). In patch 1, we introduced 2200 uninfected KS3 hosts and 240 infected ones (10% initial prevalence). Dispersal to downstream patches was allowed by opening tube connections for 3h in 2-2-3-day intervals. Prior to opening, we removed 2-ml samples from each patch to measure host density and to determine infection prevalence for patches 1, 3, 5 and 7. After 5 weeks (15 time points), the experiment ended with density and prevalence measured across all patches. We set up 24 infected landscapes (2 per parasite line); one landscape was lost due to handling error.

### Statistical analysis

For experimental data, mixed-effect models included parasite evolutionary treatment (front vs. control) as fixed factor and parasite line identity as random factor; additional factors were included as needed. Error structure was chosen according to the type of response variable. In the virulence assay, population growth and decline phases were analysed separately. For the landscape assay, local outbreak of infection prevalence was analysed separately for patches 1, 3, and 5. The rate of spatial spread was analysed by taking front patch position of host or parasite as a function of time (43), and additionally in Cox models assessing outbreak timing in downstream patches. For genomic data, we first regressed the number of nonsynonymous variants on the number of synonymous variants, in a model with evolutionary treatment as fixed factor, and COG class, COG gene and parasite line identity as random factors. This is equivalent to a genome-wide comparison of dN/dS ratios. Second, we analysed variation in the number of synonymous or nonynonymous variants, in models with evolutionary treatment and COG class as fixed factors; we additionally fitted the number of loci per COG class and parasite line as a covariate, thereby accounting for variation in coverage. This represents a multiway contingency table comparing COG diversity between evolutionary treatments. We used the R 4.2.0 statistical package “lme4”, “car”, and “survival” (71–73). Details for each analysis can be found in the respective SI sections.

### Epidemiological model

#### Single-patch dynamics

(eq. 1). In a chemostat model, asexual susceptible hosts (S) reproduce at rate b_0_, and die at rate μ_0_. Waste (W), produced at rate a and degrading at rate f, affects natality (b_1_) and mortality (μ_1_). Flow rate k reflects experimental sampling and medium renewal.

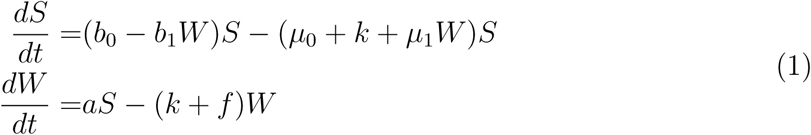

We extended this to a susceptible–infected model (eq. 2):

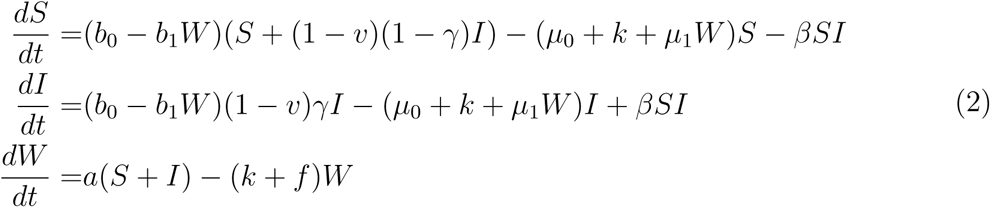

Horizontal transmission (β) occurs through direct contact between susceptible and infected hosts (I). Infection decreases host birth rate by a fraction v (fecundity virulence). Reproducing infected individuals transmit the parasite vertically with probability γ. Model parameters are listed in Table 13.1.

#### Metapopulation dynamics (*eq. 3*)

In a linear landscape, patches (i ∈ [1,N]) are connected by host dispersal (d), occurring symmetrically to adjacent patches, except at boundaries. Infection modulates dispersal by a factor λ, such that infected hosts disperse at λd. To mirror experimental dispersal episodes, dispersal rates were adjusted by a piecewise function:

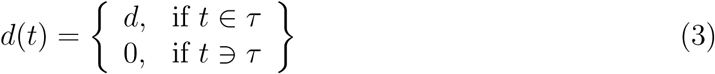

where τ denotes the time when patch connections are opened, and m a generic dispersal rate. The model was developed in Python (v3.12; Python Software Foundation).

The simulations used Gillespie’s algorithm for continuous-time systems. We varied β, v, d and λ, in 1000 different combinations, created through latin hypercube sampling (74), each replicated 3 times. Of these, 993 combinations produced spatial spread. Multiple regressions assessed the effects of β, v, d, and λ on epidemic peak size (max number of infections) and parasite arrival in patch 3 (≥5 infected hosts). Interactions between parameters were visualized with response-surface plots (R package “*rsm”* (*75*)).

## Acknowledgments

This work was funded by a grant from the Agence Nationale de la Recherche to O.K. (grant no. ANR-20-CE02-0023-01). This is publication ISEM-YYYY-XXX of the Institut des Sciences de l’Evolution – Montpellier.

## Data availability

The experimental and modelling data from this study are available at https://zenodo.org/records/20630441. The genomic data have been deposited with links to BioProject accession number PRJNA1465310 in the NCBI BioProject database (https://www.ncbi.nlm.nih.gov/bioproject/).

## Supporting Information

### Figures

**Fig. S1.**
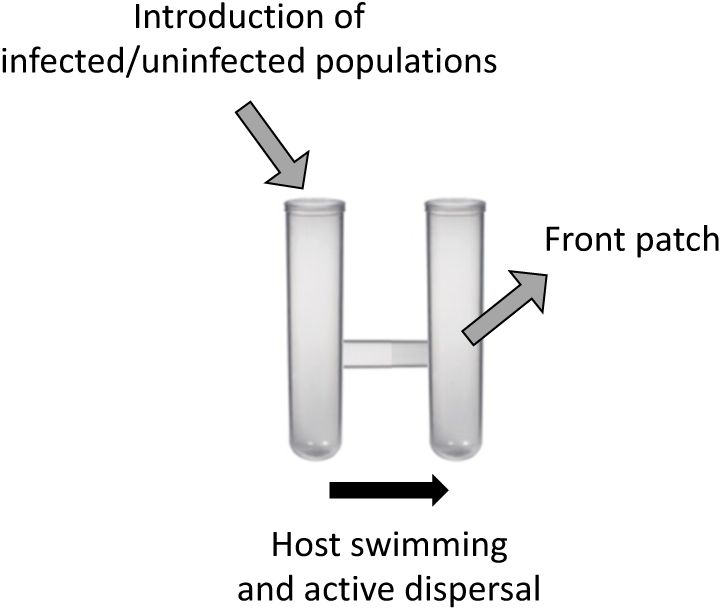
Dispersal systems used for the dispersal assays composed from two patches interconnected by 5-cm silicon tubing serving as a corridor. Once the clamp blocking the corridor was removed, the paramecia could displace from the patch in which they were introduced to the connected one (front patch) through active dispersal for 3h (black arrow). The same protocol was repeated for the wave-front and evolution-control treatment. Briefly, in the long-term experiment short dispersal episodes of 3h were alternated with weekly periods of population growth and maintenance (1 week = 1 cycle). For the wave-front, only dispersers to new front patches were propagated in the next cycle. For the evolution-control, hosts and parasites were manually transferred to new patches (matching the density of the wave-front treatment).

**Fig. S2.**
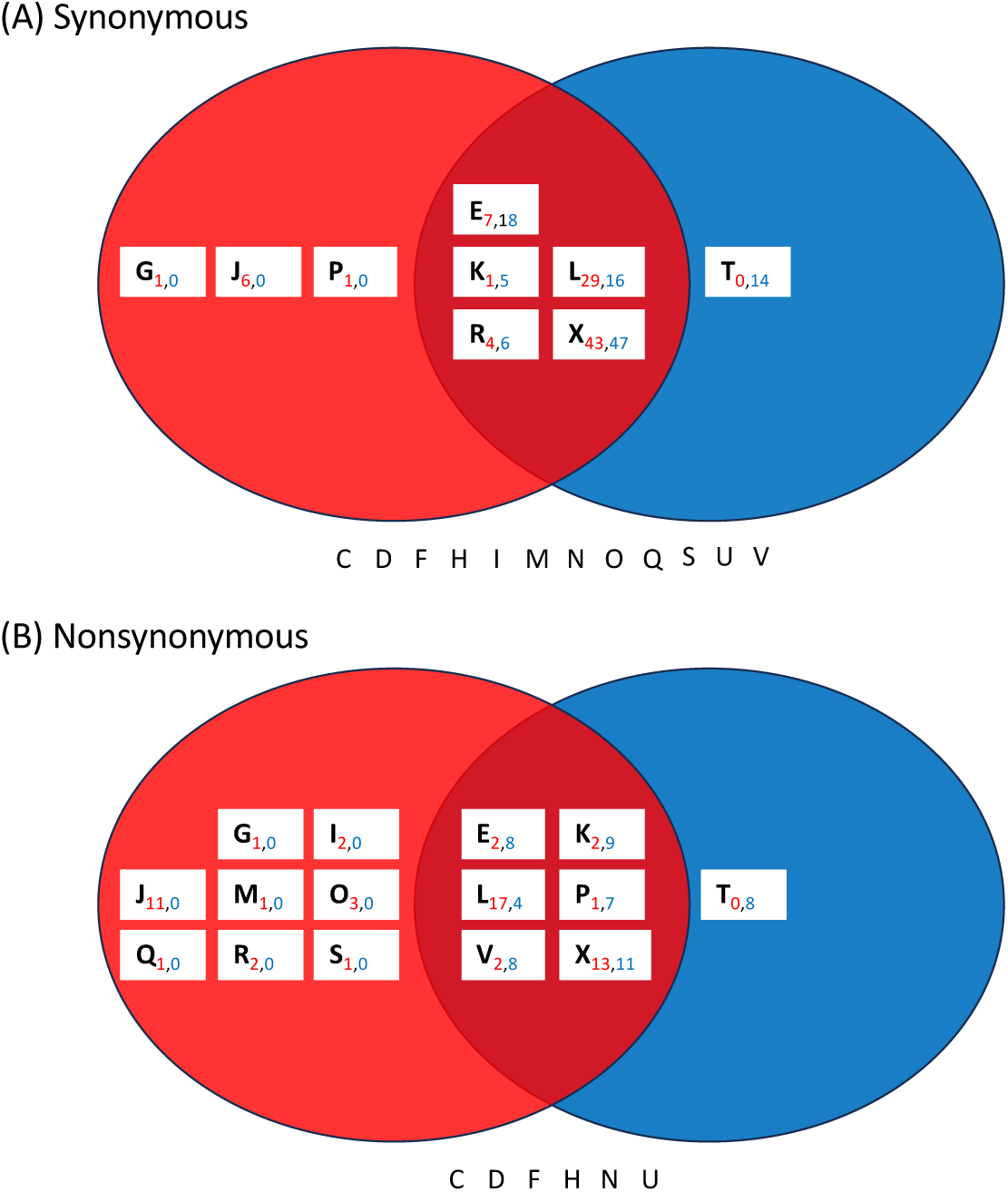
Venn diagram for (A) Synonymous and (B) Nonsynonymous variants for different COG class (white box) for front (red) and control (blue) parasites. Subscripts indicate the number of variants per parasite type. For instance, in (A) the two parasite treatments shared variants for five COG class, with E having seven front variants and 18 control variants. The letters beneath each diagrams indicate COG class for which no variants were detected.

**Fig. S3.**
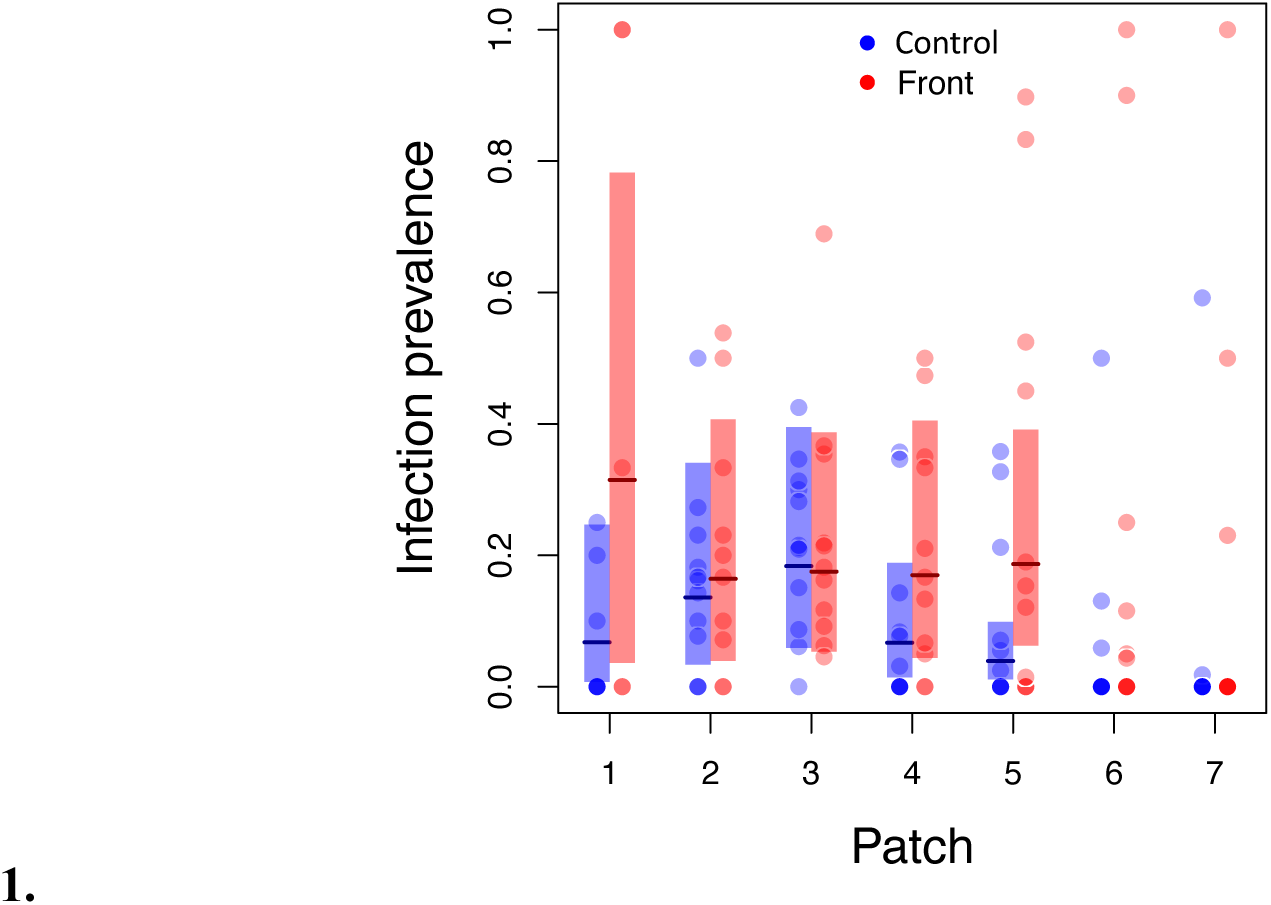
Overall infection prevalence of control (blue) and front (red) evolved parasite from patch 1 to 7 of the landscape during the last week of the epidemiological assay. Each point is an individual replicate, panels are the 95% confidence intervals of the model predictions for patch 1 to 5, which were analysed (main text). Thick lines correspond to the mean model prediction.

**Fig. S4.**
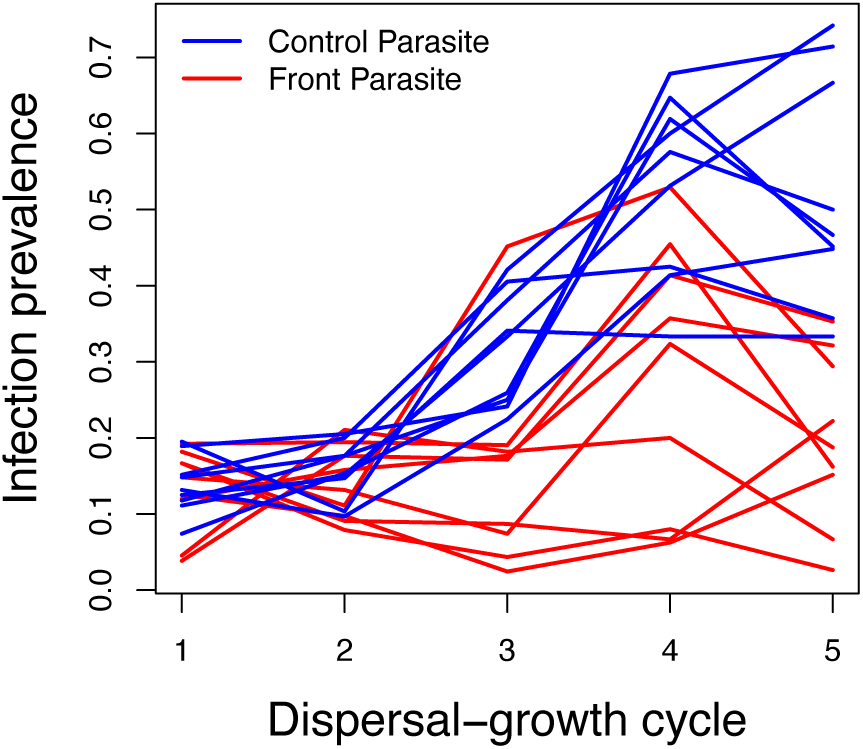
Infection prevalence over time. Infection prevalence of control (blue) and front (red) parasites for the first 5 dispersal-growth cycles of the long-term experiment. Each line represents an evolutionary replicate.

**Fig. S5.**
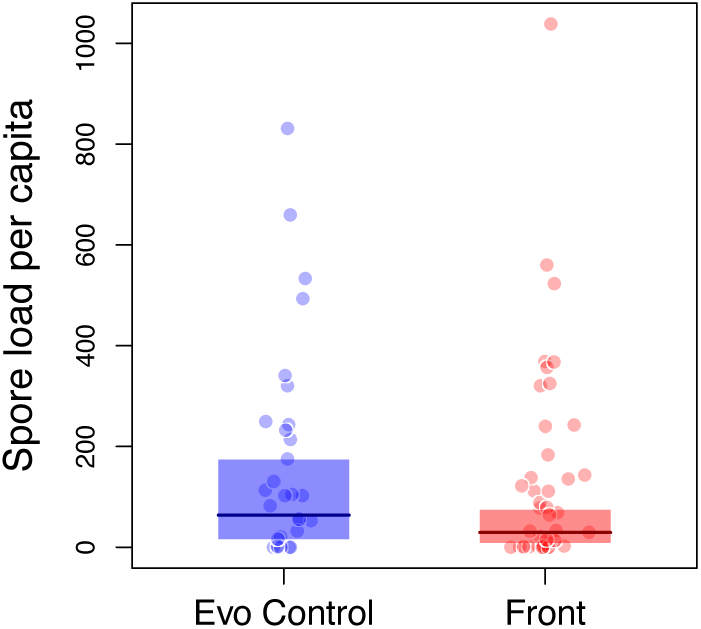
Transmission investment as spore load per capita. Each point represents the number of spores per paramecium of a replicate, thick lines and shaded areas are median and 95% CI of the model prediction. The transmission investment of control (blue) and front (red) parasites were not statistically different (χ^2^=0.83, p=0.36).

**Fig. S6.**
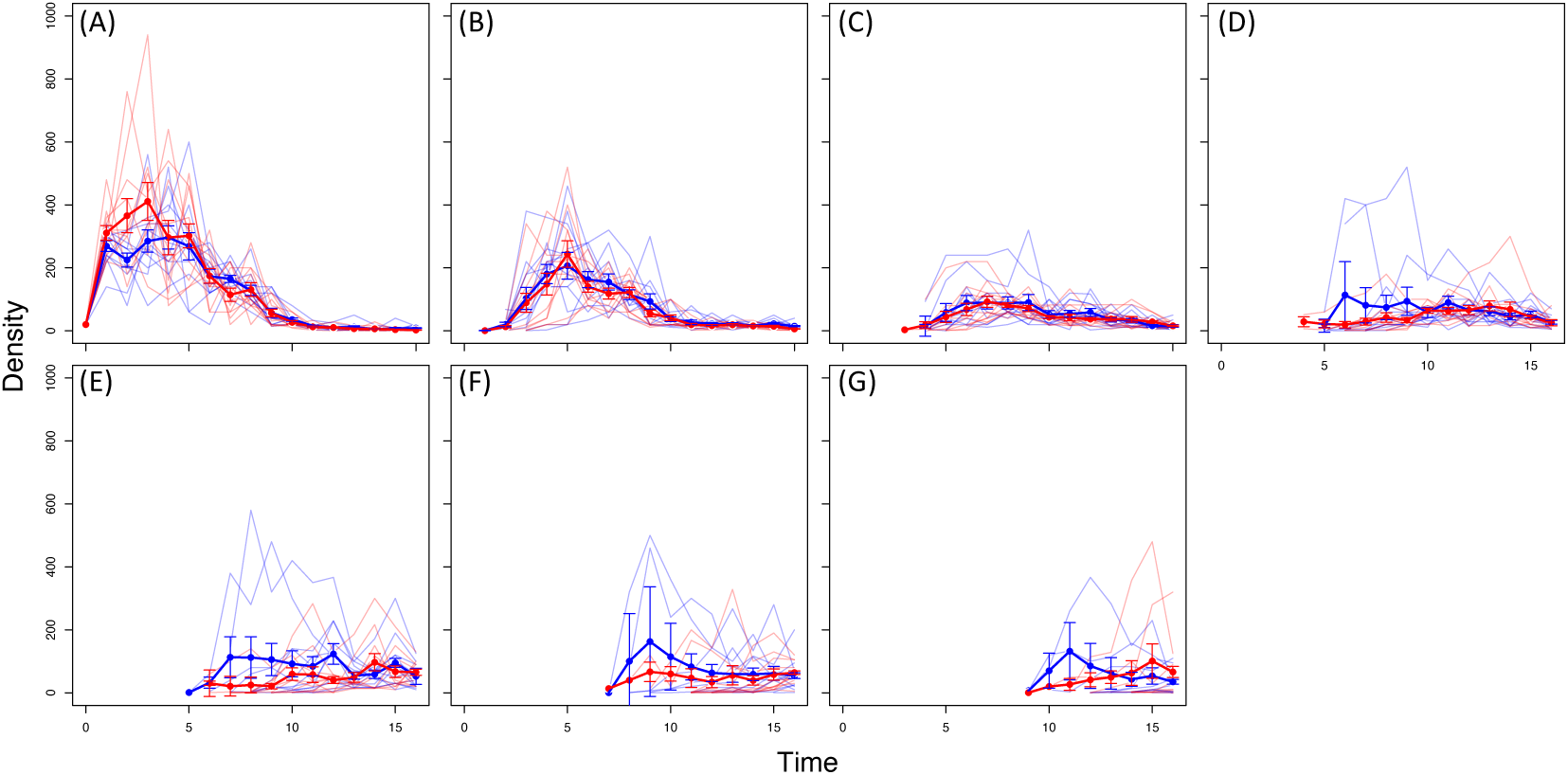
Host density over time across the landscape. (A-G) Population density from patch 1 to 7 over the sampling period (opening events), in red and blue populations exposed to front and control parasite respectively. Each point and bars represent the average density and standard error, thick lines are the average density trajectories. Shaded lines show the density trajectory of each replicate.

**Fig. S7.**
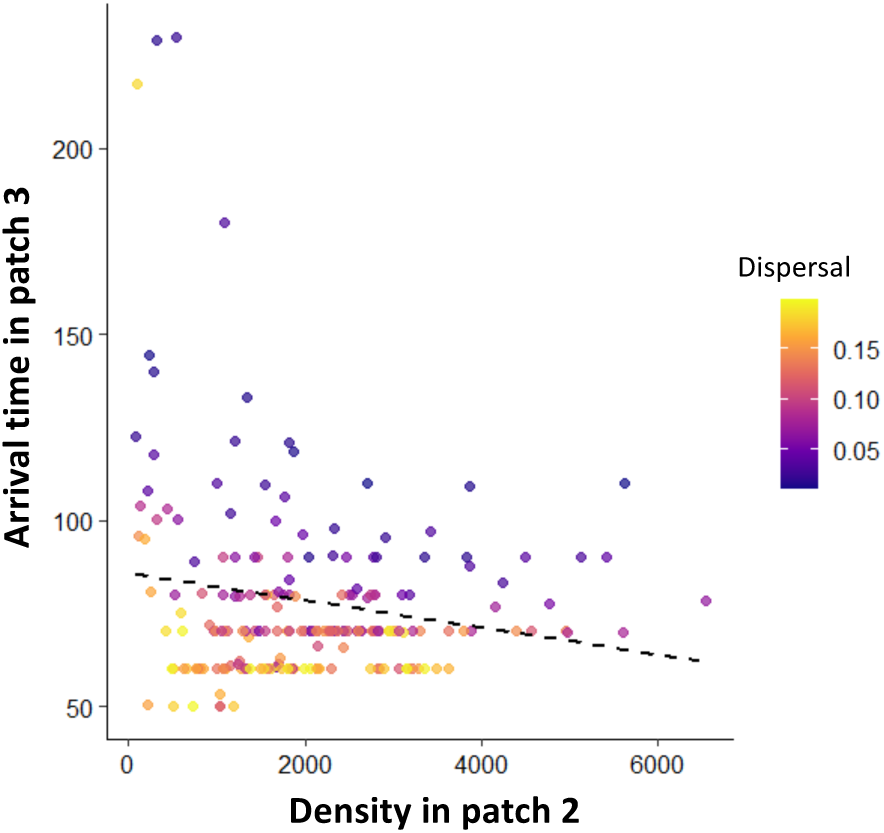
Negative relationship between parasite arrival time in patch 3 in function of the density of infected individuals in patch 2 at the previous time point. Colours indicates different dispersal levels.

### Tables

**Table S1.**
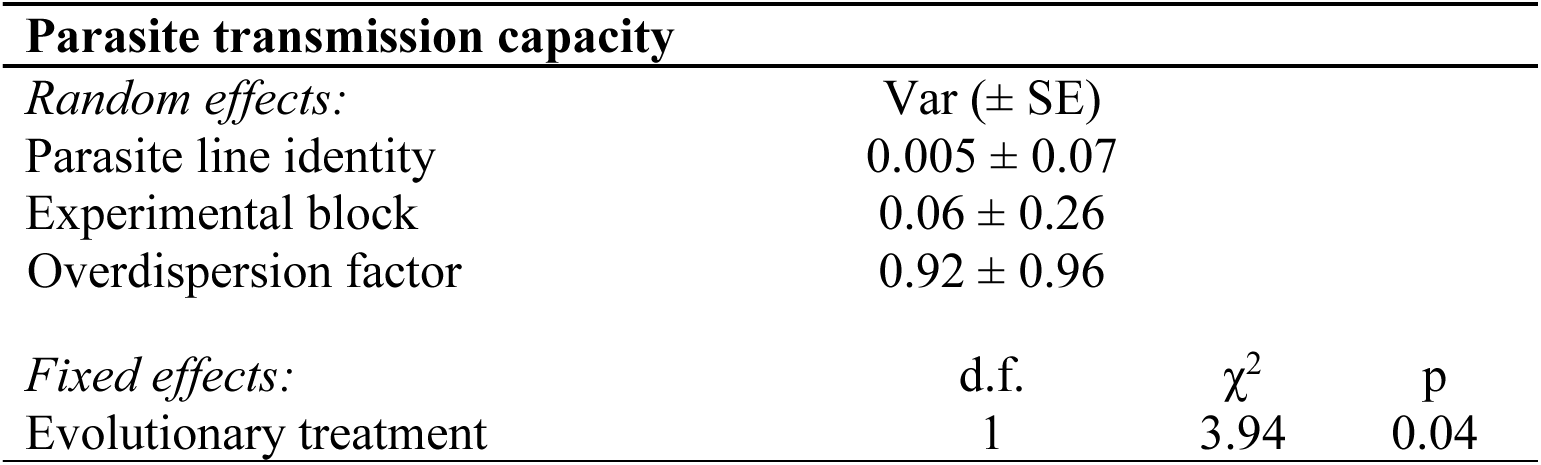
ANOVA results for GLMM model for parasite transmission capacity. For random factors, variance (Var) and standard error (SE) are provided.

**Table S2.**
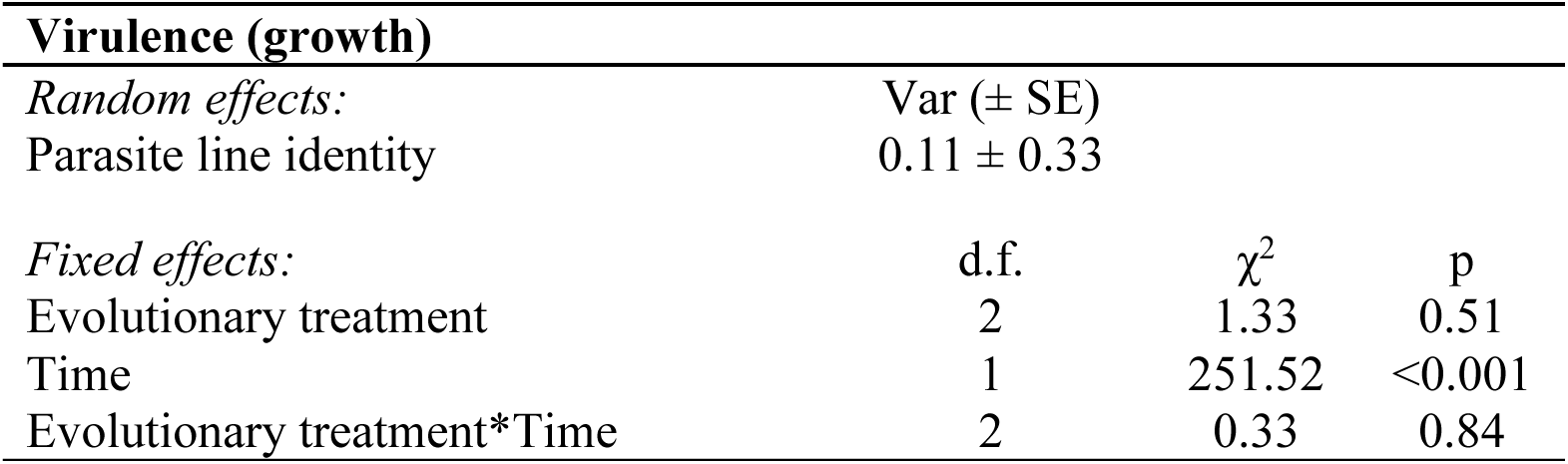
ANOVA results of LMM for parasite virulence (growth). For random factors, variance (Var) and standard error (SE) are provided. The evolutionary treatment fixed effect includes 3 factors (front parasite, control parasite, and no parasite).

**Table S3.**
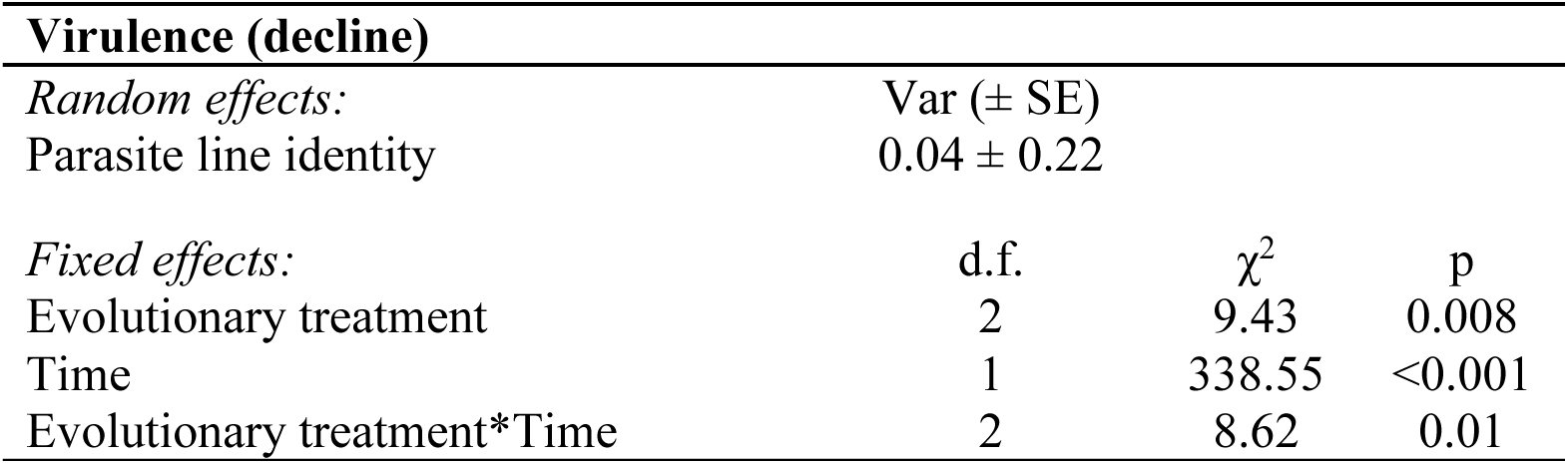
ANOVA results of LMM for parasite virulence (decline). For random factors, variance (Var) and standard error (SE) are provided. The evolutionary treatment fixed effect includes 3 factors (front parasite, control parasite, and no parasite).

**Table S4.**
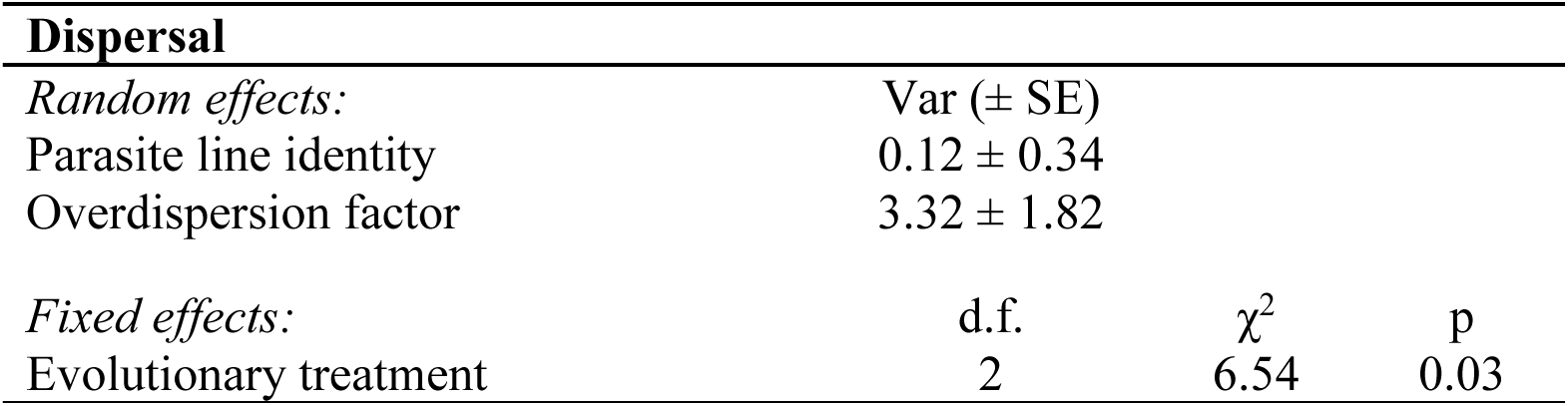
ANOVA results of GLMM for dispersal. For random factors, variance (Var) and standard error (SE) are provided. The evolutionary treatment fixed effect includes 3 factors (front parasite, control parasite, and no parasite).

**Table S5.** Total count of genetic variants per sample identified with SNIPPY. Total contigs 1161, N50 = 1645 bp; total length = 1 170 196 bp; average coverage 6.13x.

| Sample ID | Treatment | Total variants |
| --- | --- | --- |
| 4 | Front | 311 |
| 5 | Front | 445 |
| 6 | Front | 136 |
| 7 | Front | 379 |
| 8 | Front | 114 |
| 9 | Front | 167 |
| 10 | Front | 151 |
| 11 | Control | 400 |
| 12 | Control | 209 |
| 13 | Control | 340 |
| 14 | Control | 486 |
| 15 | Control | 167 |
| 16 | Control | 143 |
| 17 | Control | 104 |
| 18 | Control | 210 |
| 19 | Control | 160 |

**Table S6.** Summary of the COG categories retrieved in this study.

|  |  |
| --- | --- |
| R | General function prediction only |
| Q | Secondary metabolites biosynthesis, transport and catabolism |
| J | Translation, ribosomal structure and biogenesis |
| X | Mobilome: prophages, transposons |
| E | Amino acid transport and metabolism |
| V | Defense mechanisms |
| L | Replication, recombination and repair |
| I | Lipid transport and metabolism |
| O | Posttranslational modification, protein turnover, chaperones |
| K | Transcription |
| H | Coenzyme transport and metabolism |
| G | Carbohydrate transport and metabolism |
| T | Signal transduction mechanisms |
| N | Cell motility |
| P | Inorganic ion transport and metabolism |
| S | Function unknown |
| C | Energy production and conversion |
| D | Cell cycle control, cell division, chromosome partitioning |
| F | Nucleotide transport and metabolism |
| M | Cell wall/membrane/envelope biogenesis |
| U | Intracellular trafficking, secretion and vesicular transport |

**Table S7.**
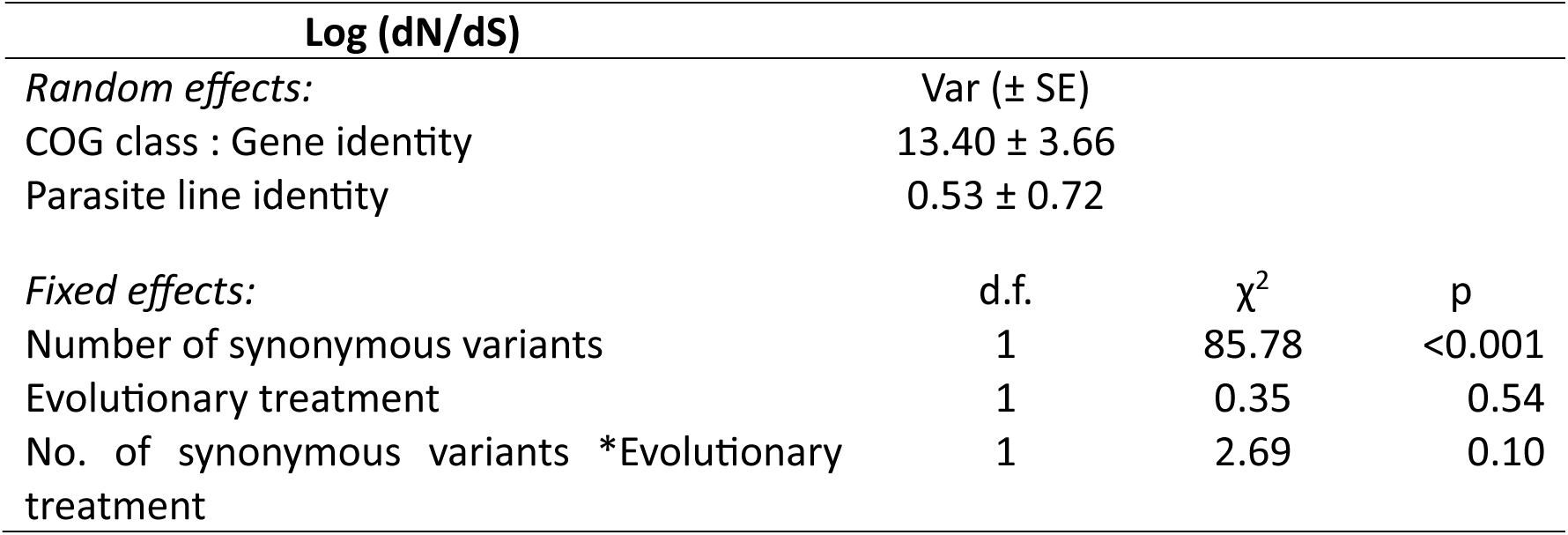
ANOVA results of GLMM model (poisson distribution) analysing variation in the number of nonsynonymous variants, as a function of the number of synonymous variants in parasite genomes from front and control treatments. Parasite line identity, COG class and COG gene identity were taken as random factors. For random factors, variance (Var) and standard error (SE) are provided. This analysis tests whether, for a given number of synonymous variants, evolutionary treatments differ in the number of nonsynonymous variants. This is equivalent to a comparison of dN/dS ratios.

**Table S8.** ANOVA results of GLM model analysing variation in the number of synonymous variants (poisson distribution), comparing parasite genomes from front and control treatments and different COG classes. This analysis is equivalent to a contingency table, testing whether variant diversity differs between evolutionary treatments (= significant COG class * evolutionary treatment interaction). Variant frequencies were combined over parasite lines and COG genes. To take into account variation in sequencing coverage (which can affect the probability of detecting variants), the model included the number of loci covered per p COG class and parasite line as a covariate.

| **Number of synonymous variants** |  |  |  |
| --- | --- | --- | --- |
| *Fixed effects:* | d.f. | $\chi^2$ | p |
| COG class | 20 | 167.59 | <0.001 |
| Evolutionary treatment | 1 | 0.29 | 0.58 |
| Number of loci covered | 1 | 0.03 | 0.84 |
| COG class * Evolutionary treatment | 20 | 19.64 | 0.48 |

**Table S9.** ANOVA results of GLM model analysing variation in the number of nonsynonymous variants (poisson distribution), comparing parasite genomes from front and control treatments and different COG classes. This analysis is equivalent to a contingency table, testing whether variant diversity differs between evolutionary treatments (= significant COG class * evolutionary treatment interaction). Variant frequencies were combined over parasite lines and COG genes. Variant frequencies were combined over parasite lines and COG genes. To take into account variation in sequencing coverage (which can affect the probability of detecting variants), the model included the number of loci covered per COG class and parasite line as a covariate.

| **Nonsynonymous variants** |  |  |  |
| --- | --- | --- | --- |
| *Fixed effects:* | d.f. | $\chi^2$ | p |
| COG class | 20 | 76.21 | <0.001 |
| Evolutionary treatment | 1 | 2.00 | 0.15 |
| Number of loci covered | 1 | 12.58 | <0.001 |
| COG class * Evolutionary treatment | 20 | 44.67 | 0.001 |

**Table S10.**
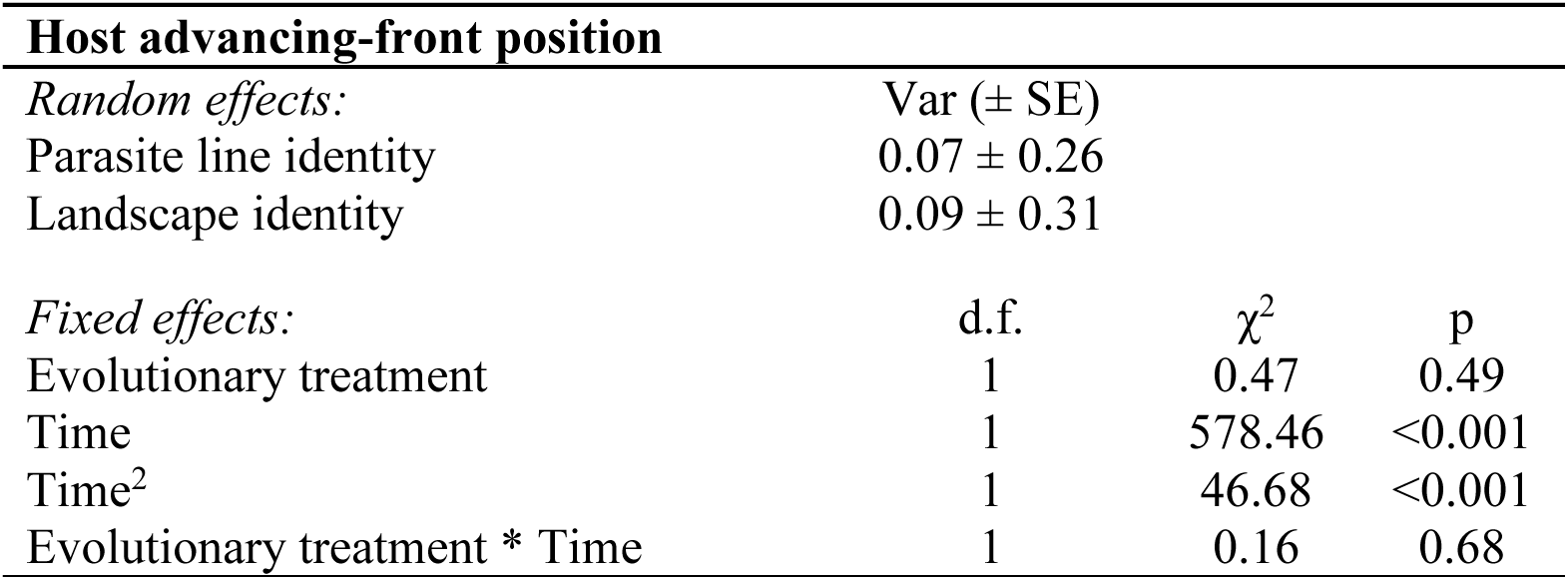
ANOVA results of LMM for host advancing-front position. For random factors, variance (Var) and standard error (SE) are provided.

**Table S11.**
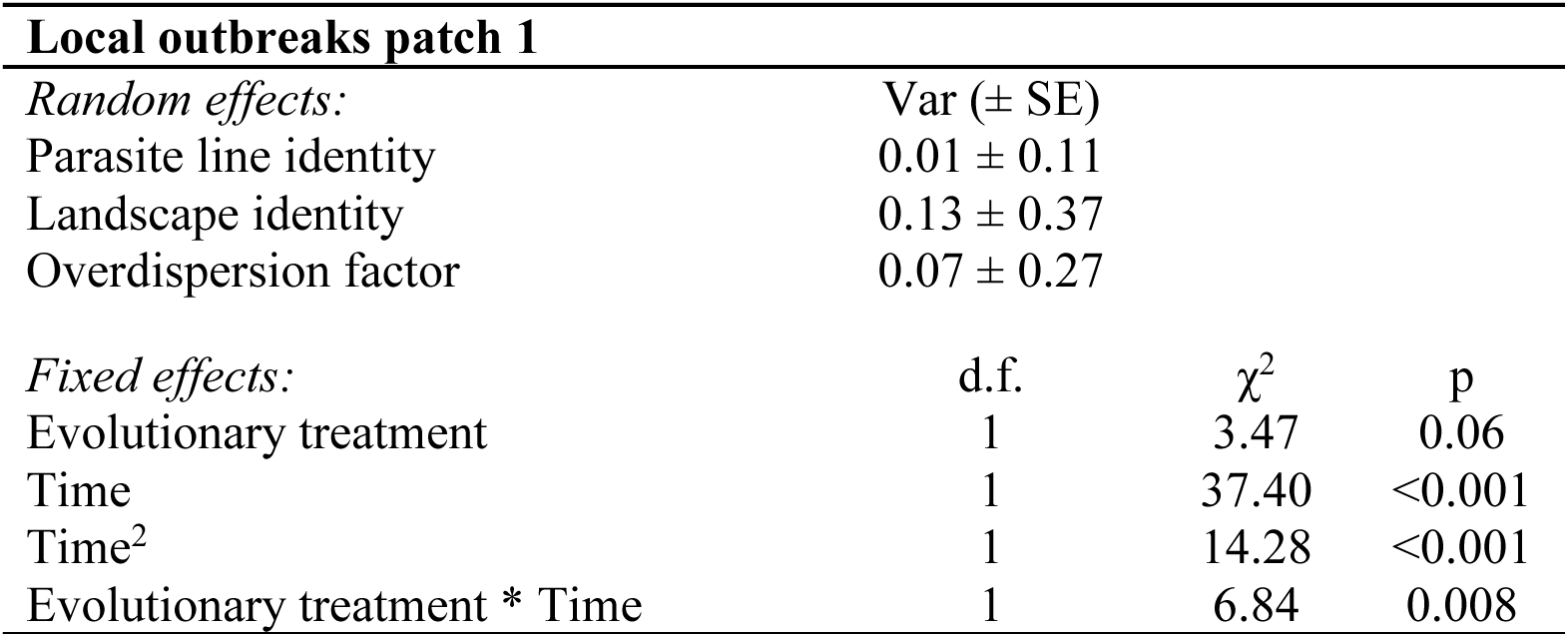
ANOVA results of LMM for local outbreaks in patch 1. For random factors, variance (Var) and standard error (SE) are provided.

**Table S12.**
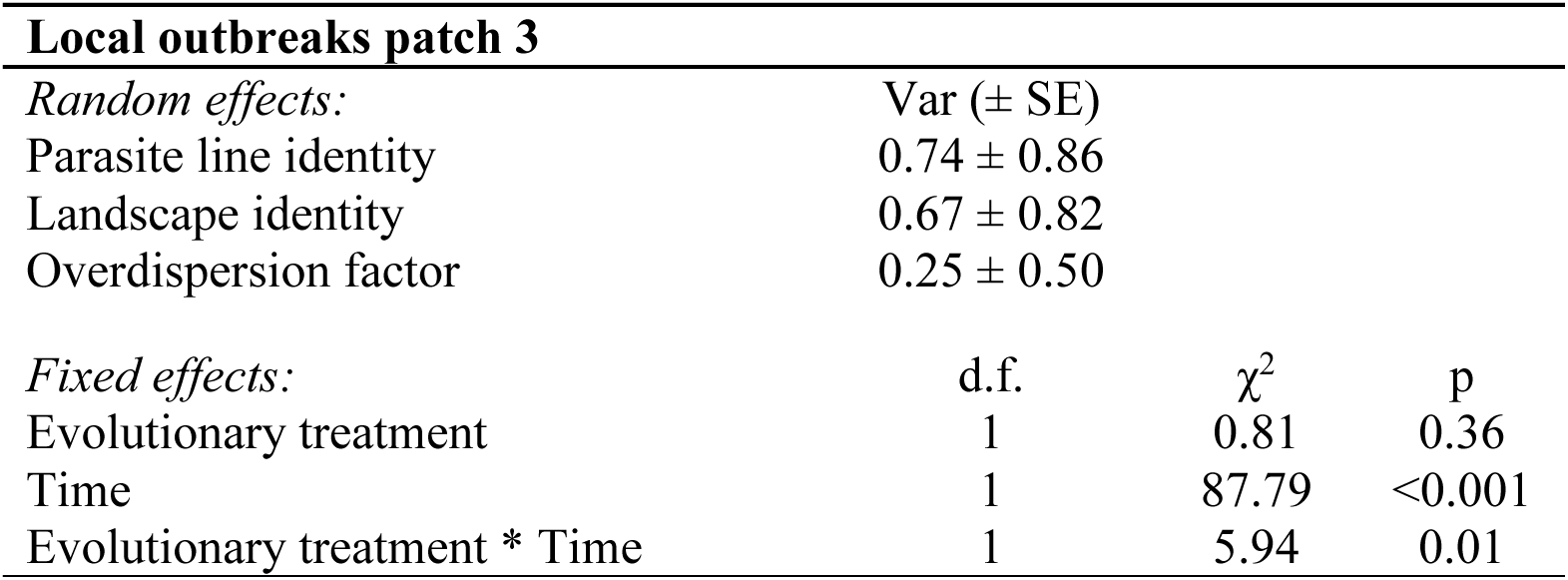
ANOVA results of LMM for local outbreaks in patch 3. For random factors, variance (Var) and standard error (SE) are provided.

**Table S13.**
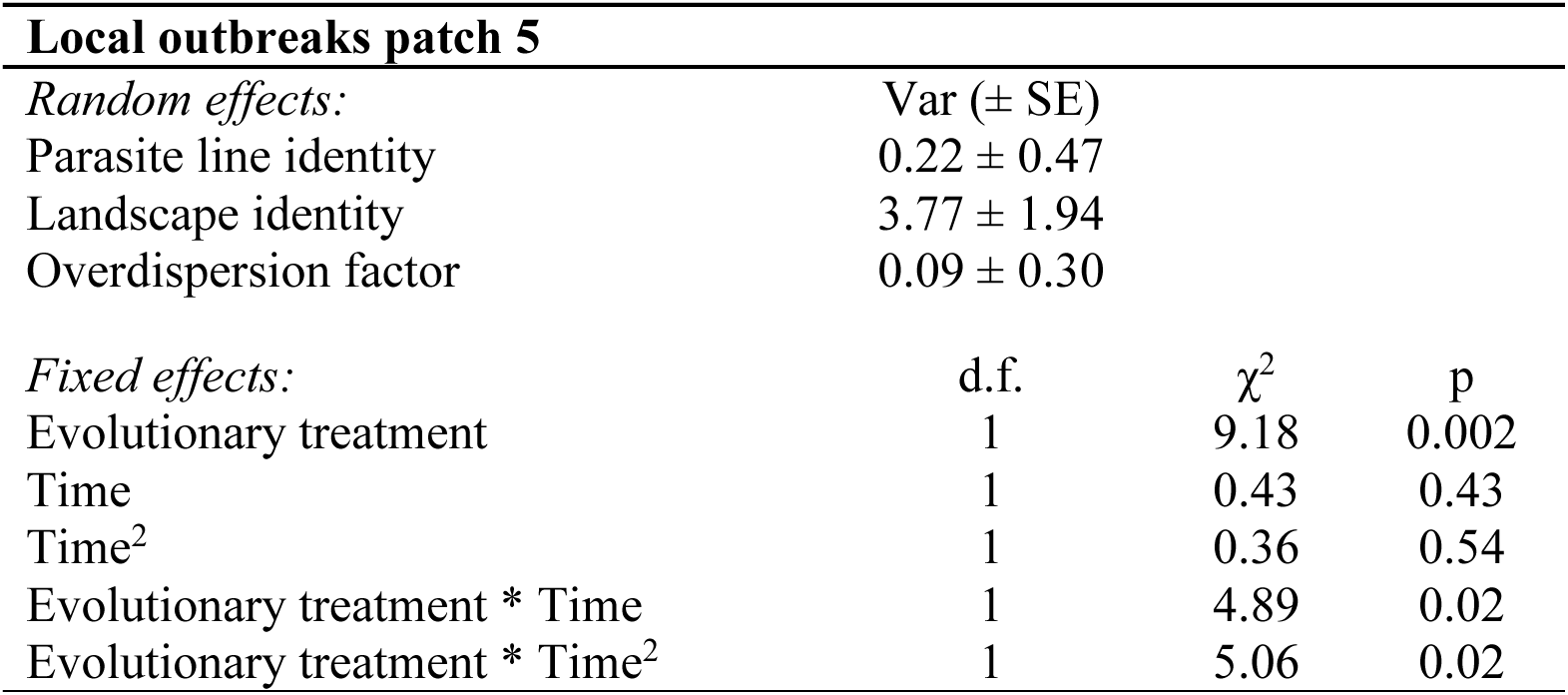
ANOVA results of LMM for local outbreaks in patch 5. For random factors, variance (Var) and standard error (SE) are provided.

**Table S14.**
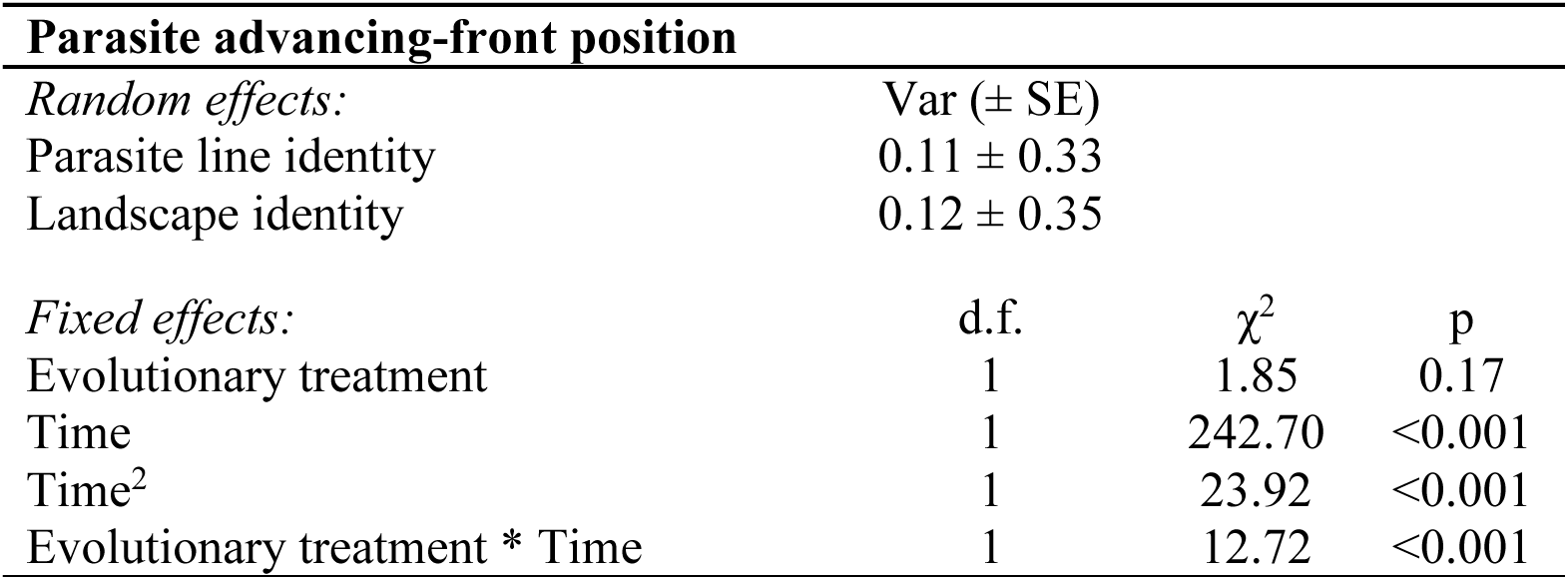
ANOVA results of LMM for parasite advancing-front position. For random factors, variance (Var) and standard error (SE) are provided.

**Table S15.** ANOVA results of Cox model for parasite time of arrival.

| **Parasite time of arrival** |  |  |  |
| --- | --- | --- | --- |
| *Fixed effects:* | d.f. | $\chi^2$ | p |
| Evolutionary treatment | 1 | 3.98 | 0.04 |
| Patch position | 1 | 20.45 | <0.001 |

**Table S16.**
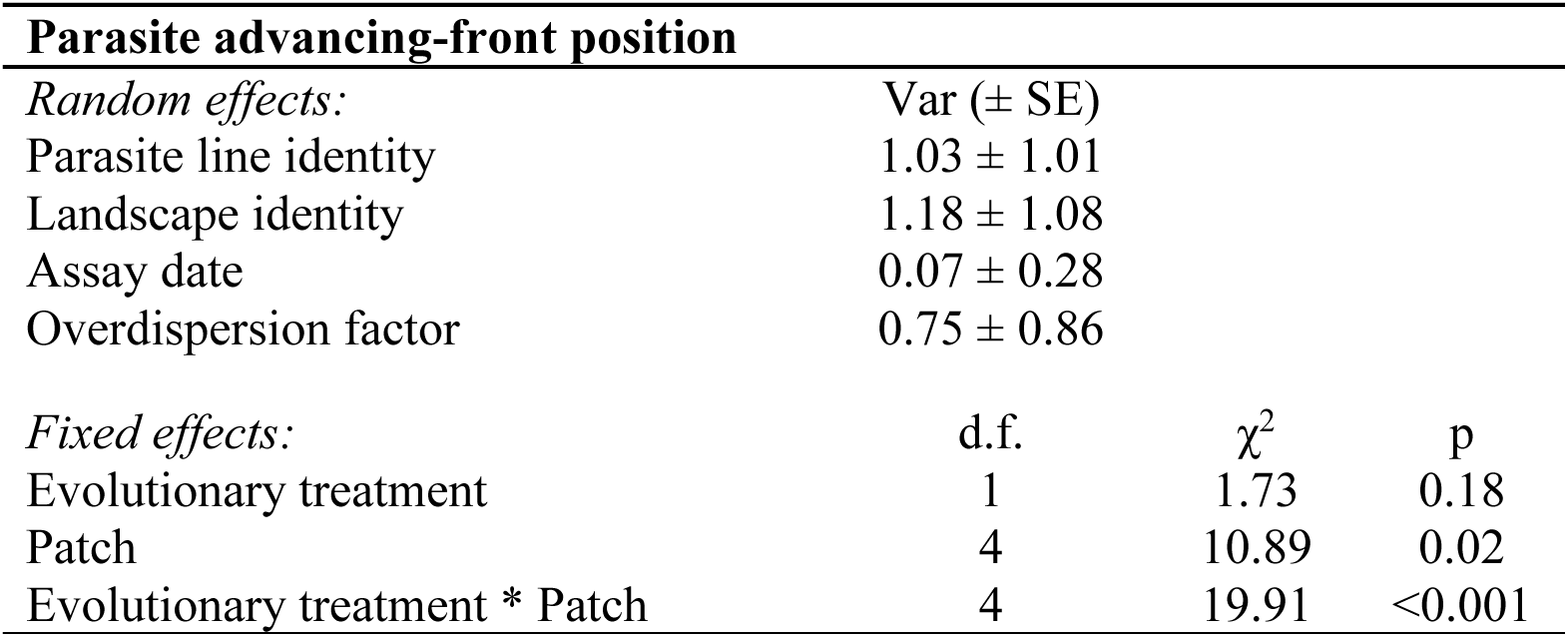
ANOVA results of GLMM model for parasite final infection prevalence across landscape. For random factors, variance (Var) and standard error (SE) are provided.

**Table S17.**
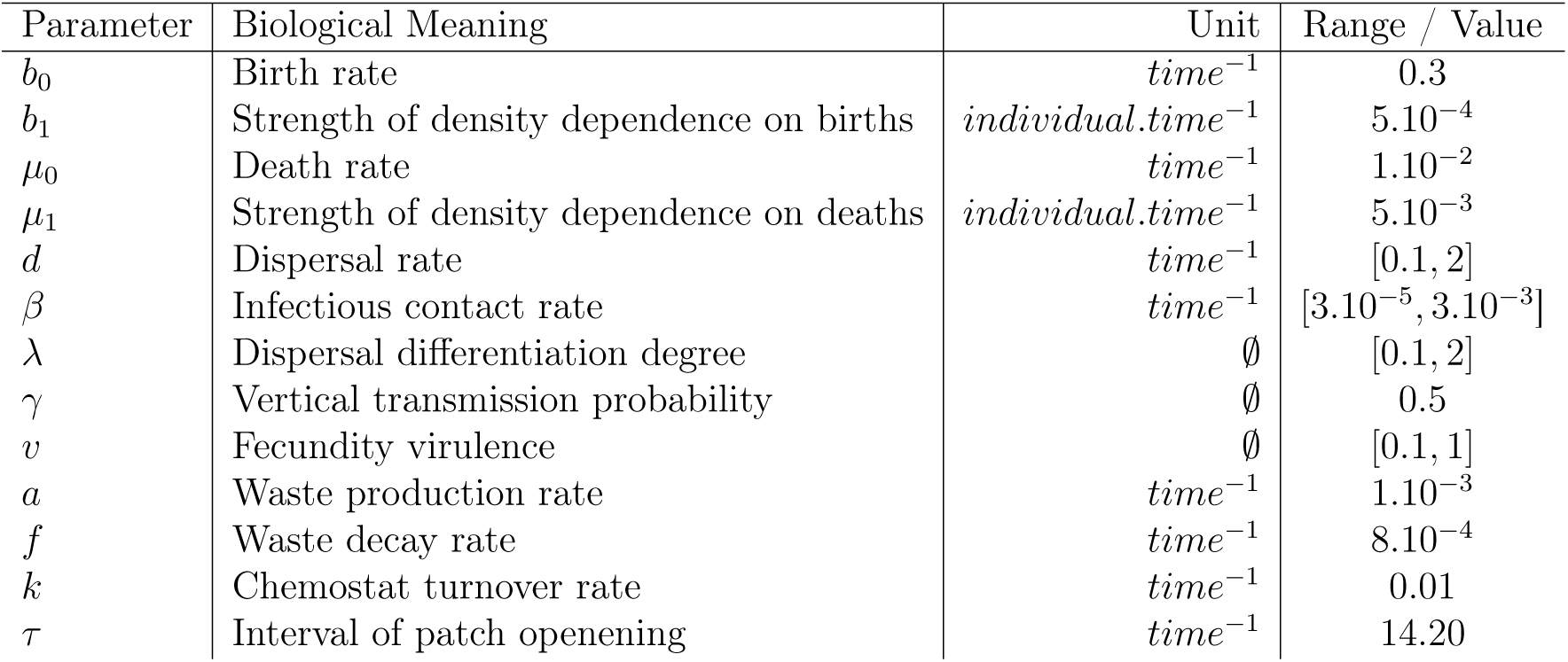
Model parameters description. Parameters with no range of variation were set constant across all simulations.

**Table S18.**
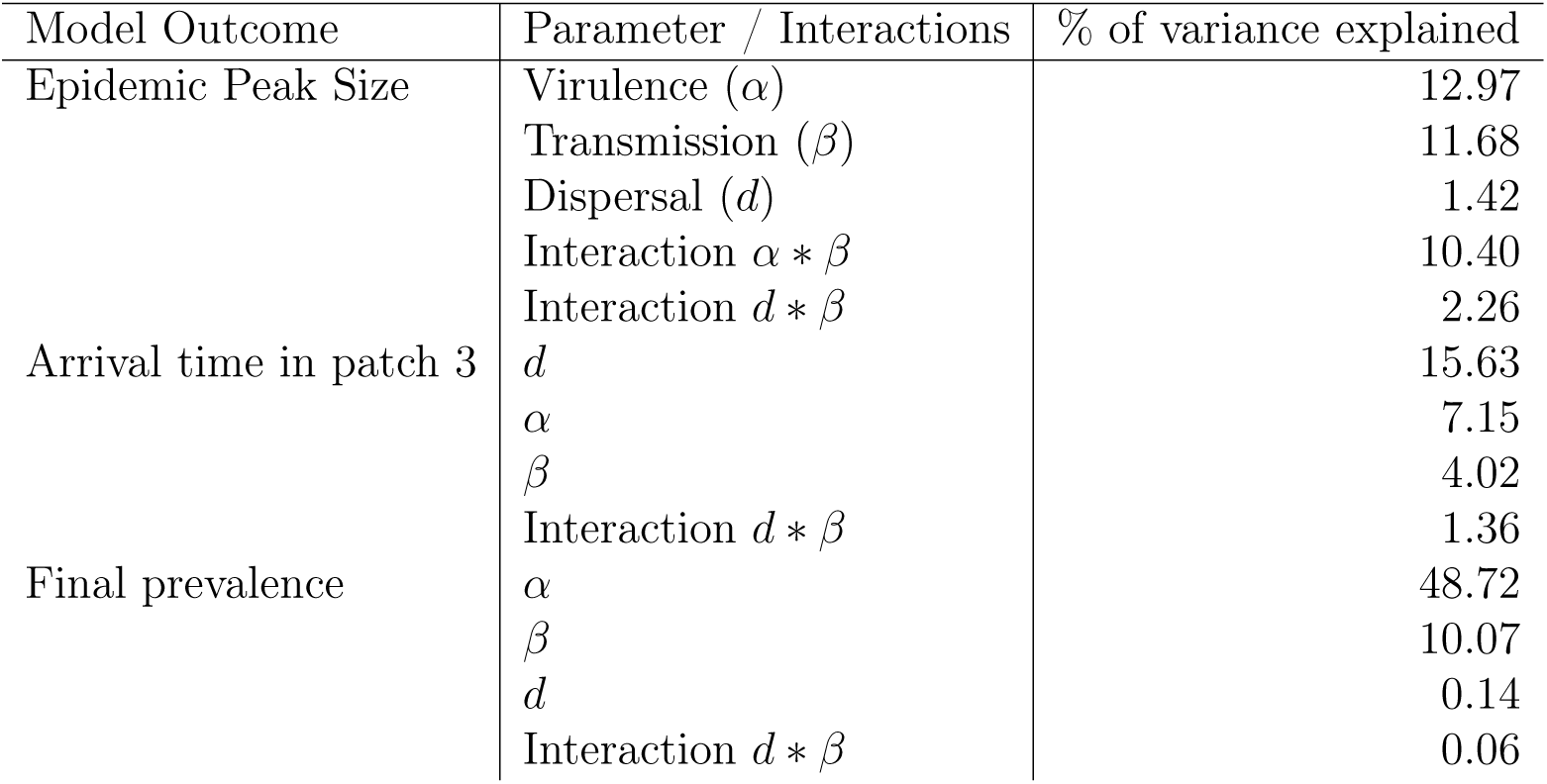
Approximated Sobol sensitivity indices for key model outcomes. The percentage of variance explained is calculated from an ANOVA on the selected sub-model derived from model averaging, using the parameters and second-order interactions identified as significant (*i.e.*, with a cumulative model-averaged weight > 0.8).

| Model Outcome | Parameter / Interactions | % of variance explained |
| --- | --- | --- |
| Epidemic Peak Size | Virulence ($\alpha$) | 12.97 |
| | Transmission ($\beta$) | 11.68 |
| | Dispersal ($d$) | 1.42 |
| | Interaction $\alpha * \beta$ | 10.40 |
| | Interaction $d * \beta$ | 2.26 |
| Arrival time in patch 3 | $d$ | 15.63 |
| | $\alpha$ | 7.15 |
| | $\beta$ | 4.02 |
| | Interaction $d * \beta$ | 1.36 |
| Final prevalence | $\alpha$ | 48.72 |
| | $\beta$ | 10.07 |
| | $d$ | 0.14 |
| | Interaction $d * \beta$ | 0.06 |

**Table S19.**
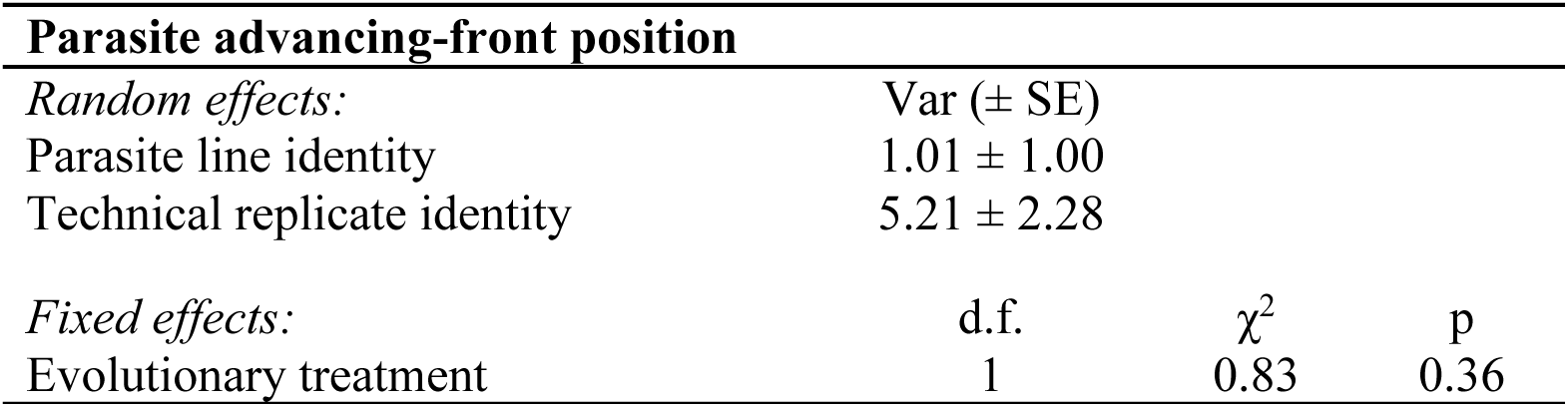
ANOVA results of GLMM model for parasite final infection prevalence across landscape. For random factors, variance (Var) and standard error (SE) are provided.

**Table S20.** ANOVA results of LMM model for the density in each patch in function of the parasite evolutionary treatment as fixed factor (control vs. front), and time as random term.

| **Density** | **d.f.** | **$\chi^2$** | **p** |
| --- | --- | --- | --- |
| *Patch 1* |  |  |  |
| Evolutionary treatment | 1 | 3.24 | 0.07 |
| *Patch 2* |  |  |  |
| Evolutionary treatment | 1 | 1.99 | 0.15 |
| *Patch 3* |  |  |  |
| Evolutionary treatment | 1 | 1.39 | 0.23 |
| *Patch 4* |  |  |  |
| Evolutionary treatment | 1 | 3.55 | 0.059 |
| *Patch 5* |  |  |  |
| Evolutionary treatment | 1 | 5.00 | 0.02 |
| *Patch 6* |  |  |  |
| Evolutionary treatment | 1 | 1.96 | 0.16 |
| *Patch 7* |  |  |  |
| Evolutionary treatment | 1 | 0.81 | 0.36 |

